# CSR Calculator: An R package and Shiny application for assigning plant ecological strategies using trait data

**DOI:** 10.1101/2025.06.06.658218

**Authors:** Teddy Gaskin, Ben Crick, Amanda P Cavanagh, Katharina Huber, Zoran Nikoloski, John N Ferguson

## Abstract

1. The competitor, stress-tolerator, ruderal (CSR) theory, first proposed by John Philip Grime, is a useful framework for understanding plant ecological strategies and predicting responses to environmental changes and pressures. However, current tools for assigning CSR strategies are limited to an Excel sheet format and have yet to be integrated into modern computational platforms that enable reproducible research.
2. We present *CSRcalculator* (https://github.com/TeddyGaskin/CSRcalculator), an open-source R package and shiny application that calculates CSR scores and assigns strategies based on user-uploaded trait data. *CSRcalculator* supports CSR assignments according to the three most prominent models: The original *soft approach*, the global StrateFy, and a morpho-physiological model.
3. The R package outputs a table including CSR scores, assigned strategies and intermediate traits for the selected model. The shiny application produces this same table alongside an interactive ternary plot to visualise the strategy distribution and an optional summary table describing the overall CSR strategy distribution, using metrics such as, the modal strategy, axis means, standard deviations, and ranges. Group-level analyses calculate the same statistics across user-defined categories.
4. We provide a worked example using our tool to assign CSR strategies to plants for an example dataset.

## 1 Introduction

Understanding how plants interact with their environment is fundamental to addressing global ecological challenges, including biodiversity conservation and climate change adaptation. The observed response of a plant to environmental challenges can be broadly categorised into competitive, stress-tolerant, and ruderal (CSR) strategies (Grime, 1974). Plants falling into the competitor category survive in relatively resource-abundant and stable environments by investing resources into rapid growth for the pre-emptive attainment of further resources, thus increasing competitive ability. In contrast, stress-tolerator plants invest in their capacity to retain resources and repair damage under stress in response to resource-limited environments. Lastly, ruderal plants survive in environments that undergo repeated destructive events, such as wildfires or human activity, by investing in their reproduction ability, allowing plants to regenerate despite these events.

The CSR framework can assist in evaluating the conditions of a plant community, for example, the prevalence of stress-tolerant species in a plant community may indicate some degree of shading; this is in contrast to a community with a high proportion of competitive species that could indicate greater light exposure (Massant, et al. 2009). Additionally, the framework can be used to predict plant responses to environmental changes and can pinpoint species that are the “winners” or “losers” due to this change. From this information we can determine whether or not human intervention is required to prevent loss of biodiversity. More recent applications of CSR employ it as a framework that can aid in the understanding of adaptive species variation as well as community level shifts in functionality. Vasseur et al. (2018) highlighted the value of the CSR framework for explaining local environmental adaptations, identifying differences in strategy classification among *Arabidopsis thaliana* accessions from distinct climates, with plants from colder regions exhibiting more stress-tolerant strategies. Zhang et al. (2024) focused on community-level strategies, and provided empirical evidence for the predicted shift from ruderal to stress-tolerant strategies during secondary succession (Grime, 1977, 2001; Grime & Pierce, 2012) demonstrating the usefulness of CSR classifications for tracking changes in plant community strategy through time.

Plant species are arranged along CSR axes based on indexing for each strategy according to specific functional traits, with classification determined by the coordinates for each species within a triangular CSR space (Hodgson et al. 1999; Table 1). These functional traits give insight into how a plant allocates its resources and are ideally representative of the extreme trade-offs for each strategy; for example, competitive species tend towards large leaf areas (LA), enabling greater resource acquisition, whilst stress-tolerant species exhibit smaller specific leaf area (SLA) coupled with high leaf dry matter content (LDMC) reflecting a thicker and tougher leaf to be more resistant to environmental stresses (Chen et al. 2019). The original CSR framework scored plant species in the C-axis according to canopy height, litter accumulation and lateral spread, and in the S-axis according to the maximum relative growth rate in the seedling phase (Grime, 1974). The R-axis, initially underdeveloped, was later refined to reflect the role of disturbance in plant strategies (Grime, 1986), leading to a seven-type classification system that included primary types (C, S, R) and intermediates (CR, SR, SC, and CSR). Further acknowledgment of 12 tertiary classes resulted in the 19 CSR categories that are currently used (Grime et al. 1988). Hodgson et al. (1999) later developed a model for CSR classification based on measurable functional traits, including canopy height (CH), lateral spread (LS), SLA, LDMC, flowering start (FS) and flowering period (FP), to provide a standardised approach for classifying plant CSR strategies. To build this model, they began by defining a set of “gold standard” CSR scores for plant species that had been judged as C, S or R based on expert opinion and ecological data: competitiveness was quantified using a ‘dominance index’ derived from the relative abundance of a species within a stable environment; stress-tolerance was based on scores on the first principal component axis (PCA) from analyses of laboratory trait data for plants grown under controlled conditions designed to simulate resource-limited environments; and ruderality was estimated from co-occurrence with monocarpic species in disturbed habitats. These gold standard values were then used as dependent variables in multiple regression analyses, with combinations of transformed functional traits selected through trial and error to maximise the correlation with each axis. The resulting regression equations form the basis of the model, allowing raw C, S, and R scores to be calculated from trait data.

**Table 1.**
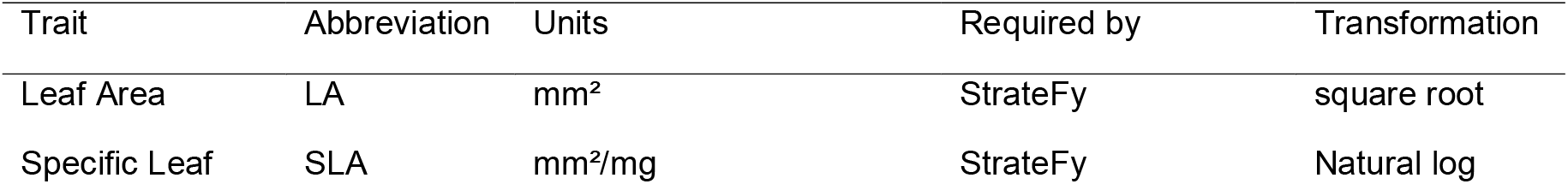
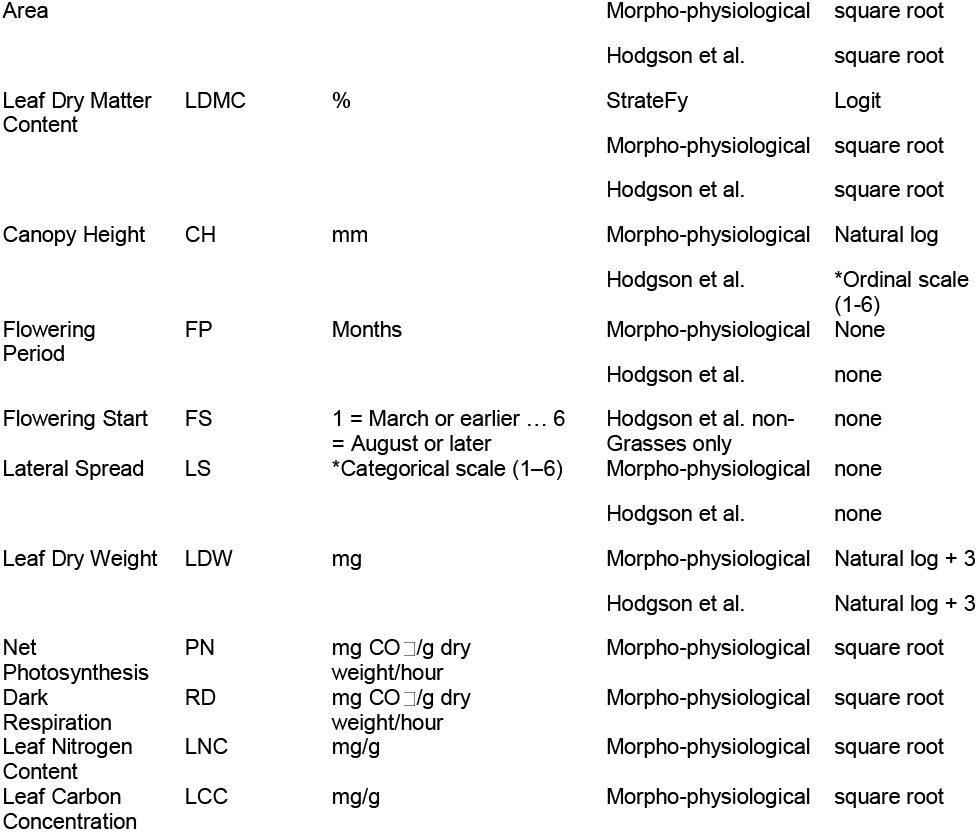
Required traits and their transformations per each CSR model. *For conversions to these scales, see Hodgson et al. (1999).

Two more recent models have further expanded the CSR framework. The StrateFy model (Pierce et al. 2017) limits the selection of functional traits to leaf traits: specific leaf area (SLA), leaf dry matter content (LDMC), and leaf area (LA), with SLA and LDMC calculated from leaf fresh weight (LFW) and leaf dry weight (LDW). For this model, equations to calculate CSR scores are derived from PCA of leaf trait data gathered from the TRY database (www.try-db.org; Kattge et al. 2011). Specifically, the first two PCA axes are used: axis 1 captured the trade-off between SLA and LDMC and represents a spectrum from competitive to stress-tolerant strategies; axis 2, influenced primarily by leaf area, helps to differentiate ruderal species. Transformed trait data were then regressed against the PCA axis that captured the greatest variation for those traits, to generate equations that estimate a species’ position in CSR space. The minimal trait set makes the model suitable for broad-scale ecological applications due to the easily accessible measurements that are ubiquitous across a variety of plant species.

The morpho-physiological model developed by Novakovskiy et al. (2016) extends the selection of functional traits with the inclusion of physiological traits such as photosynthetic capacity (PN) and dark respiration (RD) for more accurate assignment of CSR strategies, offering an alternative method for CSR classification. Similar to StrateFy, this model is also based on PCA, but here it is applied to a trait dataset including physiological and morphological traits. Axis 1 reflected most of the variation in physiological traits whilst axis 2 captured a combination of nutrient content and morphological traits, allowing species’ positions to be inferred within CSR space.

The above-described models are currently limited to Excel-based tools or require manual implementation. Both the StrateFy and Hodgson models are provided as a spreadsheet tool (Pierce et al., 2017; Hodgson et al. 1999), in contrast, the morpho-physiological model (Novakovskiy et al., 2016) is not available as a tool and must be applied manually using published equations. The reliance on spreadsheets can present some limitations, for instance: scaling for large datasets or incompatibility with modern data science tools. To address these issues, we introduce an R (R Core Team, 2024) package and Shiny (Chang et al., 2024) application designed to calculate CSR strategies for plants allowing the user to select either/or the original Hodgson’s method (Hodgson et al. 1999), the morpho-physiological model (Novakovskiy et al. 2016) and the StrateFy model (Pierce et al. 2017). By hosting these various models on one open-source platform, this application aims to improve the accessibility and reproducibility of CSR analysis for a diverse range of ecological contexts.

## 2 Methods

### 2.1 Tool structure and implementation

The *CSRcalculator* was developed in R and is available as both an R package and a Shiny web application (https://portal.bethchatto.co.uk/csr.php; local version: https://github.com/TeddyGaskin/CSRcalculator-Shiny-application). The standalone R package is available at: https://github.com/TeddyGaskin/CSRcalculator and includes three model-specific functions, strateFy(), morphoPhys(), and hodgson(), each of which can be applied to a data frame containing species information and the necessary trait data:

strateFy()

Implements the StrateFy model. Arguments:

- data: A data frame containing species-level trait values.
- calcSLA: Logical. If TRUE, calculates SLA from LA and LDW if not provided. Default is TRUE.
- calcLDMC: Logical. If TRUE, calculates LDMC from LFW and LDW if not provided. Default is TRUE.

morphoPhys()

Implements the morpho-physiological model. Arguments:

- data: A data frame containing species-level morphological and physiological trait values.
- calcLDMC: Logical. If TRUE, calculates LDMC from LFW and LDW. Default is FALSE.

hodgson()

Implements the Hodgson et al. (1999) model. Arguments:

- data: A data frame containing species-level morphological and phenological trait values.
- calcLDMC: Logical. If TRUE, calculates LDMC from LFW and LDW. Default is FALSE.
- calcSLA: Logical. If TRUE, calculates SLA from LA and LDW. Default is FALSE.
- preferNonGrasses: Logical. If TRUE, applies the non-grass equations when valid flowering start (FS) values (1–6) are present. Default is FALSE.

These functions follow the same workflow (Fig. 1) and return a data frame with additional columns for calculated intermediate traits, the CSR scores and/or percentages and the assigned strategy classes.

**Figure 1.**
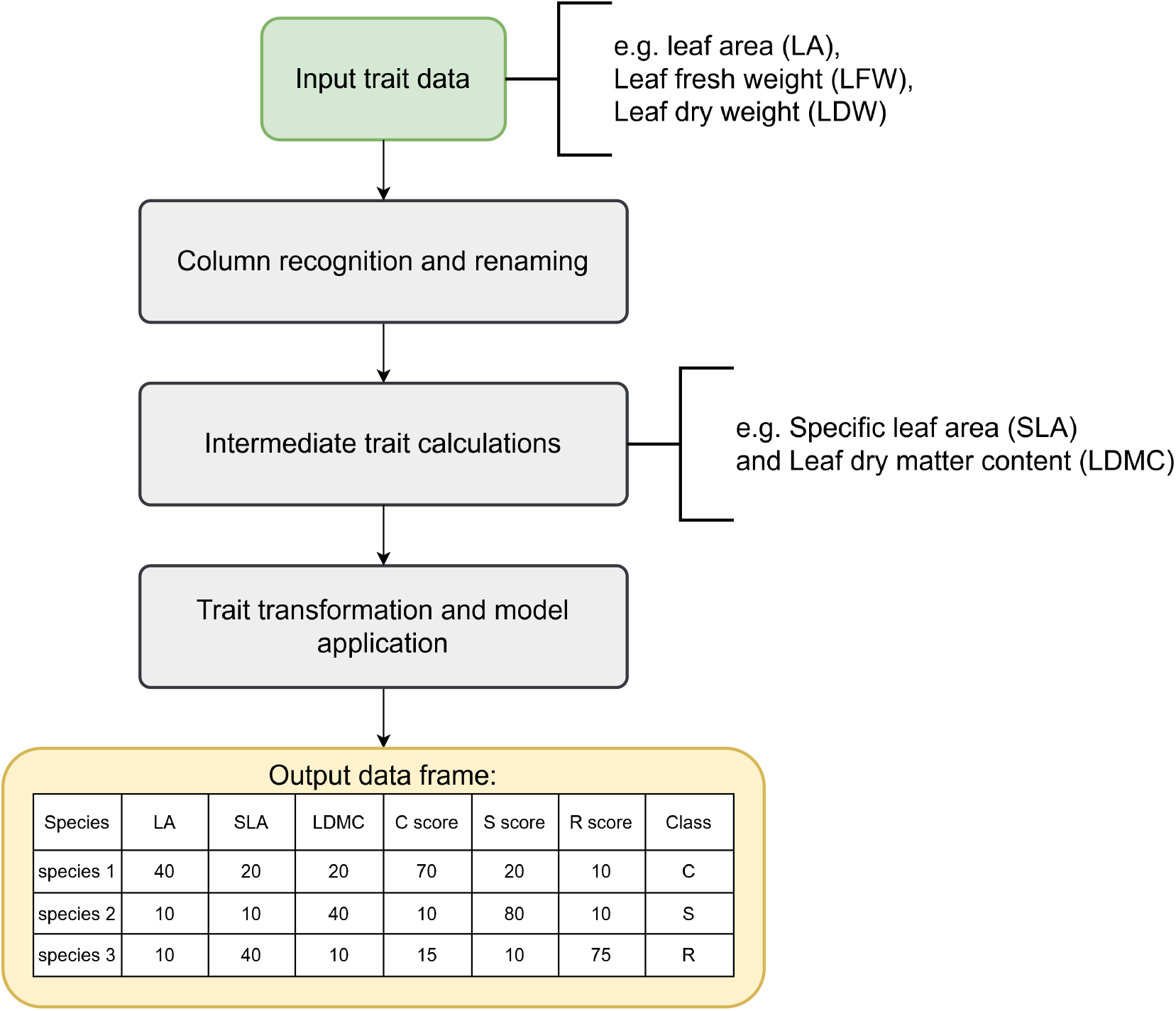
Workflow summarising the functions in the *CSRcalculator* package for assigning plant ecological strategies given trait data as input.

Alternatively, the Shiny web application hosted on: https://portal.bethchatto.co.uk/csr.php, provides a user-friendly interface for CSR analysis without requiring R experience. The user interface includes an “Introduction” panel which details usage guidance and model descriptions. The second panel, “CSR analysis”, allows the user to upload and prepare trait data. Here, users can manually assign traits to columns, select an averaging option and apply the chosen CSR model. Results are displayed in an interactive table and ternary plot. Lastly, there is an “Output” panel with options for displaying summary statistics and a degree of table and plot customisation (Fig. 2).

**Figure 2.**
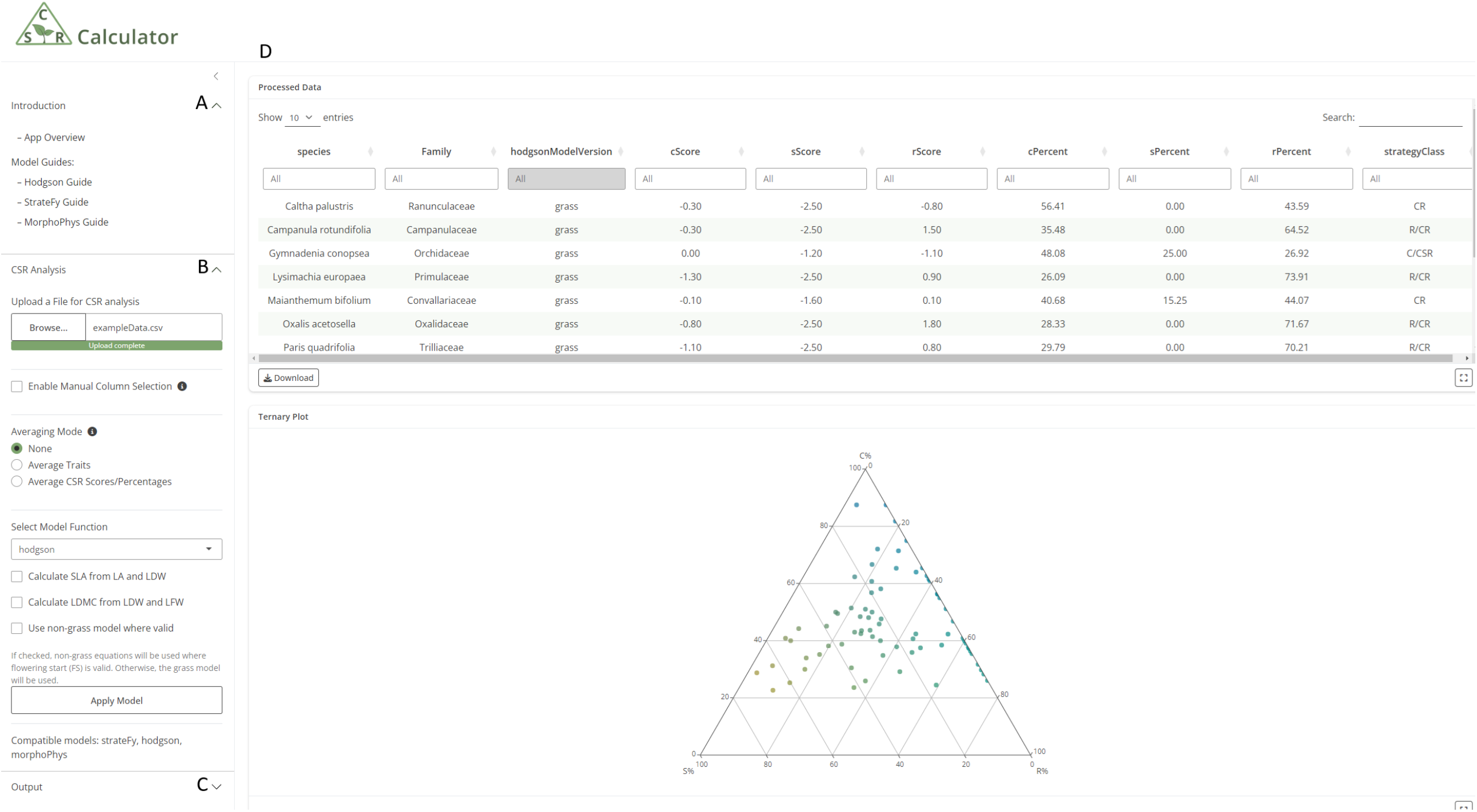
Overview of the CSR Calculator shiny application. (A) model guides, (B) analysis panel with column selection, averaging options and model application, (C) output panel with table and plot options, (D) main panel presenting results.

### 2.2 CSR model calculations

CSR model calculations are based on Pierce et al. (2017), Hodgson et al. (1999), and Novakovskiy et al. (2016). Full calculations are provided in the supporting data (Supporting Methods S1). Each model requires a specific set of traits with transformations applied prior to the model calculations (Table 1). Where applicable, intermediate traits (e.g. for non-succulents, LDMC = [LDW * 100) / LFW] or SLA = LA / LDW) are computed automatically if not provided. Trait transformations are applied internally prior to model-specific CSR calculations, following the methods described in the original publications.

In the case of the StrateFy model, the original excel tool calculates SLA and LDMC from LA, LFW and LDW, this is to account for succulent species when calculating LDMC values before their use in the model; a succulence index is calculated for each species, if this value exceeds the threshold of 5, the LDMC calculation is adjusted to prevent distortion from high water content leaves. After trait transformations, these values are used in principal component analysis (PCA)-based equations to gather axis scores for each CSR strategy. These scores are normalized and converted to percentages for strategy assignment based on the minimum Euclidean distance between CSR percentages of 19 predefined categories.

The morpho-physiological model, similarly, relies on two PCA-derived equations. Each PCA axis represents a weighted linear combination of all transformed input traits. The resulting PCA1 and PCA2 values are then projected onto CSR axes by measuring the position of each species along lines that connect the origin point of PCA1 and 2 to the centroid of known C, S and R groups. The species position along these lines is considered their new coordinates in CSR space, or their C, S and R scores. A strategy class is assigned based on the minimum Euclidean distance to one of 19 predefined CSR categories, following the same classification method as the other models, the morphoPhys() function will then convert these scores into percentages to be plotted onto a ternary plot to visualize these results, any negative percentages are clipped to zero and the difference is distributed evenly across the other strategies.

The Hodgson model is not based on PCA and instead calculates each axis score directly from transformed trait values using two sets of algebraic equations, one for grasses and one for non-grasses, they differ in trait inclusion as well as coefficients, affecting the contribution of each trait to each axis score. Although the authors do not provide rationale for this approach, it is likely due to shared morphological and phenological traits that necessitate different trait weightings to ensure accurate strategy assignment between these two groups. A C score is derived from squared values of CH and LS; the non-grass version additionally includes squared log-transformed LDW. S score is calculated from combining the squared transformations of LDMC, SLA, CH and LS. Again, the non-grasses version diverges with the inclusion of the squared transformed LDW. R score is calculated by applying a linear equation to FP, log-transformed LDW, and square root-transformed SLA, the non-grass version includes FS as well. The resulting axis scores are scaled to a coordinate space bounded between –2.5 and 2.5, then classifying each species by identifying the nearest predefined strategy.

### 2.3 Comparisons with original tools

To confirm consistency, we present one-to-one comparisons (Fig 3) between the output of the CSR calculator strateFy() function and the original StrateFy Excel tool (Pierce et al. 2017). Similar comparisons were performed for the Hodgson et al. (1999) Excel tool and the CSR scores reported in the supplementary data of Novakovskiy et al. (2021), as shown in supporting Figure 1. These comparisons are based upon the same supplementary data (Novakovskiy et al. 2021).

**Figure 3.**
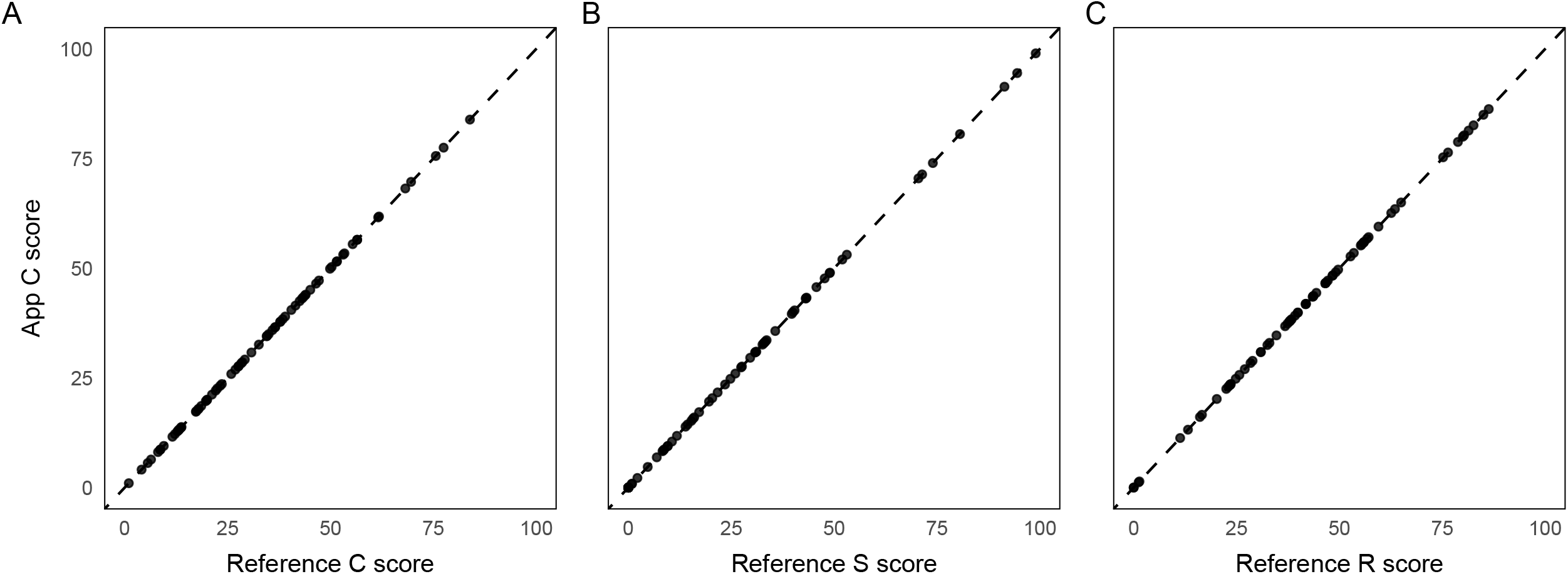
Comparison between C (A), S (B) and R (C) score outputs from the StrateFy excel tool (Pierce et al. 2017) and the *CSRcalculator* tools (Application) using supplementary trait data from Novakovskiy et al, 2021. Each panel shows a 1 to 1 assignment for each strategy, between the *CSRcalculator* application and the StrateFy excel tool.

### 2.4 Example

Here we present example results which can be reproduced by working through the available trait data using the *CSRcalculator* application (Table S1; Novakovskiy et al. 2021).

The Novakovskiy et al. (2021) dataset was uploaded to the *CSRcalculator* interface, and each model was applied using their default parameters; intermediate traits (SLA and LDMC) were calculated for StrateFy, but the morpho-physiological and Hodgson models used directly provided values. Each model produced a table displaying CSR scores and strategy assignment (consolidated in supporting Table S1), as well as a table of summary statistics (Table 2), these results were visualized through ternary plots (Fig 4).

**Table 2.** Summary table of CSR analysis for each model including the most common strategy types, the averages and standard deviation of CSR scores. Analysis based on trait data included in the supplementary data of Novakovskiy et al. (2021).

| Statistic | Hodgson<br>(grasses) | Hodgson (non-grasses) | StrateF<br>y | morpho-physiological |
| --- | --- | --- | --- | --- |
| Modal Strategy | CR | CR | CR | C/CSR |
| Mean C% | 45.98 | 51.92 | 33.18 | 33.00 |
| Mean S% | 20.25 | 10.17 | 24.55 | 35.48 |
| Mean R% | 33.77 | 37.90 | 42.27 | 31.52 |
| SD C% | 15.08 | 15.01 | 19.22 | 17.22 |
| SD S% | 19.51 | 17.92 | 25.34 | 18.03 |
| SD R% | 18.47 | 21.09 | 20.89 | 17.39 |
| Min C% | 22.67 | 26.09 | 1.02 | 0.00 |
| Min S% | 0.00 | 0.00 | 0.00 | 0.89 |
| Min R% | 2.74 | 0.00 | 0.00 | 0.00 |
| Max C% | 87.50 | 88.46 | 83.93 | 64.66 |
| Max S% | 68.49 | 68.18 | 98.98 | 74.15 |
| Max R% | 73.91 | 72.22 | 86.34 | 81.00 |

**Figure 4.**
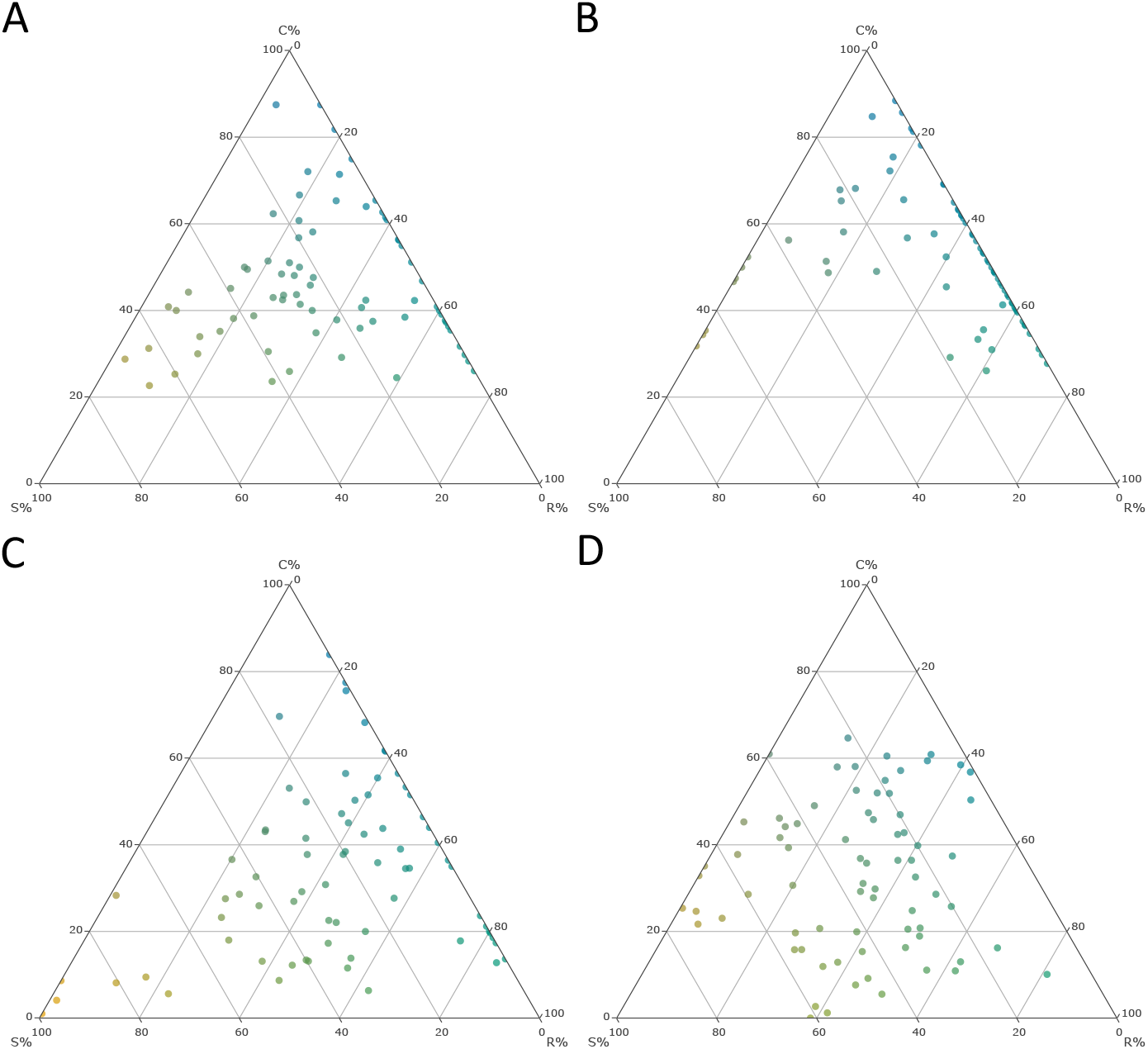
A comparison of ternary plot outputs for each CSR model using the Novakovksiy, et al. (2021) supplementary data set. (A) Hodgson (grasses) (B) Hodgson (non-grasses) (C) StrateFy and (D) morpho-physiological. Plot points are drawn towards each corner based on their CSR scores, competitive species to the top, stress-tolerators to the bottom left and ruderals to the bottom right. Point colours are blended based on their CSR scores, blue is competitive, orange is stress-tolerating and green is ruderal.

There is a large shift of species that have been classified as predominantly stress-tolerating by the morpho-physiological model towards the CR region where StrateFy and Hodgson models both score these species a 0% on the S axis. For example, *Paris quadrifolia* is classified S/CSR by the morpho-physiological model (C 11.92 %, S 52.81 %, R 35.27 %), driven largely by its relatively slow dark-respiration rate (RD = 0.42 mg CO_ g^1^ h^1^; Supporting Table 1). Both StrateFy and Hodgson classify it R/CR with S = 0 %. Likewise, *Primula matthioli* is classified S/CSR (C 15.79 %, S 56.55 %, R 27.66 %) by the morpho-physiological model, likely due to a low net photosynthetic rate indicating a slower metabolism, a characteristic of other stress-tolerating species (PN = 1.93 mg CO_ g^1^ h^1^; Supporting Table 1), yet the other models place it in CR (S = 0 %). Notably, when switching from the grass to the non-grass version of the Hodgson model, many species shift towards more extreme positions, mainly towards the CS or CR boundaries within the CSR triangle, highlighting the importance of incorporating the time of year of floral transitioning in defining CSR of non-grasses.

## 3 Model comparison and usage guidance

### 3.1 Comparing CSR models

The example data set shows distinct difference in strategy assignment across the three models for several species. This divergence stems from the differences in calculations and selection of input traits between these models. The StrateFy model is the most simplified regarding trait selection. It relies solely on three accessible morphological traits making it particularly useful for global analyses where data availability may be limited. However, this minimalistic trait selection may not capture the full spectrum of plant strategies. In contrast, the morpho-physiological model requires a minimum of 10 traits, incorporating gas exchange and nutrient content variables. The inclusion of these traits allows the model to better capture physiological trade-offs and therefore may be able to differentiate species with similar morphological traits but differing physiological adaptations. It could be argued that the CSR strategies obtained by the morpho-physiological model showed better agreement with the basic principles underlying Grime’s theory, particularly for accurately identifying stress-tolerant species. The morpho-physiological model appears to identify-stress tolerant species the other models miss, namely for species adapted to shade, cold climates and nutrient-poor soils (e.g. *Maianthemum bifolium*), rather than drought alone where the S axis seems to be primarily driven by LDMC in the StrateFy model. Both models derive their CSR scores from PCA, but the interpretation of the PCA axes differ in important ways. For StrateFy, PCA axis 1 represents the trade-offs between competitive and stress-tolerant species, where SLA is loaded positively onto this axis and LDMC, negatively. PCA axis 2 is primarily influenced by LA which helps identify potentially ruderal species. The morpho-physiological model on the other hand has axis 1 capturing most of the physiological performance differences and axis 2 is a combination of nutrient content and morphological traits. To quantify the degree of agreement between axis percentage scores for the StrateFy and morpho-physiological models, we calculated Spearman’s rank correlation coefficient (ρ) for each axis using R v4.3.2 (R Core Team, 2023). Agreement was highest for the Ruderal (R) axis (ρ = 0.68), with lower correlations for the Competitive (C) and Stress-tolerant (S) axes (ρ = 0.28 and 0.26, respectively).

The Hodgson model represents an intermediate as it relies on morphological and phenological traits but is missing physiological variables. This becomes evident in the example dataset, where the Hodgson model often agrees with StrateFy in assigning a zero score on the stress-tolerant axis to species such as *Oxalis acetosella* (Supporting Table S2) despite the morpho-physiological model classifying it as predominantly stress-tolerant. The Hodgson model is further distinct in that the CSR scores are not calculated from PCA derived equations, it instead uses regression equations developed by multiple regression analysis linking a set of “gold standard” benchmarks for each strategy to the selected traits for the model. These benchmarks were defined separately for each axis (Hodgson et al. 1999): competitiveness was estimated using a dominance index based on field survey data, stress-tolerance from PCA scores of trait data linked to nutrient stress response, and ruderality from the frequency of which a species co-occurs with short-lived plants in disturbed habitats. Notably, these standards were based primarily on British flora, which may influence the model’s applicability to an extent. Additionally, the Hodgson model includes two distinct sets of equations, one for grasses and one for non-grasses, and switching between them can substantially alter strategy assignments. Consequently, careful consideration should be given to which Hodgson model is selected based on the incorporated study species and trait data available.

### 3.2 Recommendations for use

The choice of CSR model should be guided by the scale of the study, the availability of trait data, and the level of ecological resolution that is required. Each model offers possible advantages and disadvantages depending on the context of their use:

The StrateFy model is well suited to large-scale ecological studies, where the data coverage for diverse species across regions may be a concern. Its minimal trait requirements make it a practical option for studies such as global or meta-analyses, situations where not all datasets will have a complete set of variables to use the other two models. This accessibility comes at the cost of the model’s ability to differentiate between finer ecological differences that are not morphologically based and so the determined life strategy is not always in agreement with Grime’s original theory. despite this, StrateFy remains a convenient and efficient entry point for CSR analysis.

The morpho-physiological model is the preferred choice when data availability is not a limiting factor. The inclusion of physiological traits affords better representation of plant function and resource use trade-offs between strategies. Therefore, analysis with this model is especially useful for considering subtle physiological differences that may indicate a certain ecological strategy. Although gas exchange and nutrient content measurements are timely and costly to gather, the greater resolving power of this model is a worthy trade-off for carrying out more extensive data preparation.

The Hodgson model remains useful as a benchmark. It formed the basis of early CSR classification tools and while it lacks physiological input and depends on certain morphological traits that may not be readily available, it offers a consistent reference point for validating alternative CSR models.

## 4 Conclusion

This study introduces two tools that streamline analysis of ecological strategies via the CSR framework: the open-source *CSRcalculator* R package and the accompanying shiny web application. These tools offer a more accessible and reproducible approach to CSR analysis. We have integrated the most prominent CSR models into one platform allowing researchers to choose the most appropriate model for their dataset and scale of study. In addition to facilitating current methods, the R package, and by extension shiny application, provide a foundation for future model development. New CSR models can be readily integrated into both tools, supporting the advancement and standardisation of CSR analysis in modern plant ecology.

## Supporting information

Excel sheet containing table 1 (trait data) and table 2 (CSRcalculator output).

R code text files used in the CSRcalculator R package and shiny application.

Equations and methods used for each model function in the CSRcalculator R package.

Comparison of CSR scores produced by model sources and the CSRcalculator application.

## Author Contributions

JF designed the research. TG developed the application and wrote the paper. JF, BC, AC, KH, and ZN supervised the research and discussed and approved the final version of the manuscript.

## Acknowledgements

The authors acknowledge the advice of Jo Water and Julia Boulton in the development of the application. This research was funded by ARIES DTP studentship awarded to TG.

## Data Availability Statement

All trait data used in this study were sourced from previously published materials (Novakovskiy et al. 2021). An adapted data set from this source is included in the CSR Calculator package. No new data were collected.

## Supporting information

Supporting Figure S1: Comparison of CSR scores produced by model sources and the *CSRcalculator* application.

Supporting Table S1 and 2: Excel sheet containing table 1 (trait data) and table 2 (*CSRcalculator* output).

Supporting Methods S1: Equations and methods used for each model function in the *CSRcalculator* R package.

Supporting Code: R code text files used in the *CSRcalculator* R package and shiny application.

## References

Chang, W., Cheng, J., Allaire, J., Sievert, C., Schloerke, B., Xie, Y., Allen, J., McPherson, J., Dipert, A. & Borges, B. (2024) shiny: Web application framework for R. R package version 1.10.0. Available at: https://CRAN.R-project.org/package=shiny [Accessed 3 May 2025].

Chen, H., Huang, Y., He, K., Qi, Y., Li, E., Jiang, Z., Sheng, Z. & Li, X. (2019) Temporal intraspecific trait variability drives responses of functional diversity to interannual aridity variation in grasslands. Ecology and Evolution, 9, 5731–5742.

Grime, J.P. (1974) Vegetation classification by reference to strategies. Nature, 250, 26–31.

Grime, J.P. (1977) Evidence for the existence of three primary strategies in plants and its relevance to ecological and evolutionary theory. The American Naturalist, 111(982), 1169–1194.

Grime, J.P. (1986) Manipulation of plant species and communities. In: Bradshaw, A.D., Goode, D.A. & Thorpe, E. (eds.) Ecology and design in landscape. Oxford: Blackwell Scientific, pp. 175–194.

Grime, J.P., Hodgson, J.G. & Hunt, R. (1988) Comparative plant ecology: A functional approach to common British species. Dordrecht: Springer. Available at: 10.1007/978-94-017-1094-7

Grime, J.P. (2001) Plant strategies, vegetation processes, and ecosystem properties. 2nd edn. Chichester: John Wiley & Sons.

Grime, J.P. & Pierce, S. (2012) The evolutionary strategies that shape ecosystems. Chichester: Wiley-Blackwell.

Hodgson, J.G., Wilson, P.J., Hunt, R., Grime, J.P. & Thompson, K. (1999) Allocating C-S-R plant functional types: A soft approach to a hard problem. Oikos, 85, 282–294.

Kattge, J., Díaz, S., Lavorel, S., Prentice, I.C., Leadley, P., Bönisch, G., et al. (2011) TRY – a global database of plant traits. Global Change Biology, 17, 2905–2935.

Massant, W., Godefroid, S. & Koedam, N. (2009) Clustering of plant life strategies on meso-scale. Plant Ecology, 205, 47–56.

Novakovskiy, A.B., Maslova, S.P., Dalke, I.V. & Dubrovskiy, Y.A. (2016) Patterns of allocation CSR plant functional types in Northern Europe. International Journal of Ecology, 2016, 1323614.

Novakovskiy, A.B., Dubrovskiy, Y.A., Dalke, I.V. & Maslova, S.P. (2021) Plant CSR types in the north: Comparing the morphological and morpho-physiological approaches. Physiology and Molecular Biology of Plants, 27, 665–673.

Pierce, S., Negreiros, D., Cerabolini, B.E.L., Kattge, J., Díaz, S., Kleyer, M., Shipley, B., Wright, S.J., Soudzilovskaia, N.A., Onipchenko, V.G., van Bodegom, P.M., Frenette-Dussault, C., Weiher, E., Pinho, B.X., Cornelissen, J.H.C., Grime, J.P., Thompson, K., Hunt, R., Wilson, P.J., Buffa, G., Nyakunga, O.C., Reich, P.B., Caccianiga, M., Mangili, F., Ceriani, R.M., Luzzaro, A., Brusa, G., Siefert, A., Barbosa, N.P.U., Chapin III, F.S., Cornwell, W.K., Fang, J., Fernandes, G.W., Garnier, E., Le Stradic, S., Peñuelas, J., Melo, F.P.L., Slaviero, A., Tabarelli, M. & Tampucci, D. (2017) A global method for calculating plant CSR ecological strategies applied across biomes world-wide. Functional Ecology, 31, 444–457.

R Core Team (2024) R: A language and environment for statistical computing. Vienna, Austria: R Foundation for Statistical Computing. Available at: https://www.R-project.org/ [Accessed 3 May 2025].

Vasseur, F., Sartori, K., Baron, E., Fort, F., Kazakou, E., Segrestin, J., Garnier, E., Vile, D. & Violle, C. (2018) Climate as a driver of adaptive variations in ecological strategies in Arabidopsis thaliana. Annals of Botany, 122, 935–945.

Zhang, Y.-s., Meiners, S.J., Meng, Y., Yao, Q., Guo, K.Guo, W.-Y. & Li, S.-p. (2024) Temporal dynamics of Grime’s CSR strategies in plant communities during 60 years of succession. Ecology Letters, 27, e14446.

