## Supplementary material for "CSR Calculator: An R package and Shiny application for assigning plant ecological strategies using trait data": R code text files used in the CSRcalculator R package and shiny application.: csrServer.txt

# CSR Server: Setup, Reactives, and Helpers

# Load required helper functions
source("csrFunctions.R", local = TRUE)

# Define main server function
csrServer <- function(input, output, session) {
# defining all of the reactives
  # Core Data: Uploaded and Processed Versions
  originalData             <- reactiveVal(NULL)  # Uploaded data set (raw)
  renamedData              <- reactiveVal(NULL)  # After manual or automatic column renaming
  traitAveragedData        <- reactiveVal(NULL)  # After averaging selected traits by group
  processedModelData       <- reactiveVal(NULL)  # After applying selected CSR model
  csrAveragedData          <- reactiveVal(NULL)  # After averaging CSR scores by group

  # App State: Workflow and Analysis Status
  modelApplied             <- reactiveVal(FALSE)    # Whether a model has been run
  csrAveragingConfirmed    <- reactiveVal(FALSE)    # Whether CSR score averaging is confirmed
  manualSelectionActive    <- reactiveVal(FALSE)    # Whether manual column selection is active
  averagingActive          <- reactiveVal(FALSE)    # Whether trait averaging is active
  compatibleModels         <- reactiveVal(NULL)     # List of compatible models based on traits

  # UI State: Page Content and Display Controls
  mainContentState         <- reactiveVal("intro")  # Active section of the UI (intro, guide, analysis)
  expandedCard             <- reactiveVal(NULL)     # Which output card (table/plot) is expanded
  expandedPlot             <- reactiveVal(FALSE)    # Whether the ternary plot is expanded
  redrawPlot               <- reactiveVal(0)        # Counter to trigger plot redraw
  tablePageLength          <- reactiveVal(10)       # Number of rows per page in data tables

  # Helper Functions:

  # Return the current active dataset depending on user actions
  getActiveInputData <- function() {
    # function checks for what data set is not null (!is.null = is not null) in order of what data set should be used
    if (!is.null(traitAveragedData())) {
      return(traitAveragedData())
    } else if (manualSelectionActive()) {
      return(renamedData())
    } else {
      return(originalData())
    }
  }

  # Remove duplicated columns from a dataset
  removeDuplicateColumns <- function(df) {
    # function to apply to the current data frame, [] to select, first argument empty as we don't select rows
    # second arg checks that column names have no duplicates, if this is false and there are duplicates, the selected columns are dropped
    df[, !duplicated(names(df)), drop = FALSE]
  }

  # Reset all app states to initial defaults
  resetAppState <- function() {
    renamedData(NULL)
    traitAveragedData(NULL)
    processedModelData(NULL)
    csrAveragedData(NULL)
    modelApplied(FALSE)
    csrAveragingConfirmed(FALSE)
    expandedCard(NULL)
    expandedPlot(FALSE)
    redrawPlot(0)
  }

  # Update compatible models based on available trait columns
  updateCompatibleModels <- function(data) {
    compatibleModels(checkCompatibleModels(data))
  }

  # Get the list of required traits for a given CSR model
  getRequiredTraits <- function(modelName) {
    # switch to select list of required traits based on selected model name
    switch(modelName,
           "strateFy"    = c("species", "LA", "SLA", "LDMC", "LFW", "LDW"),
           "hodgson"     = c("species", "CH", "LDMC", "FP", "LS", "LDW", "SLA", "LFW", "LA"),
           "morphoPhys"  = c("species", "CH", "LDMC", "FP", "LS", "LDW", "PN", "RD", "LNC", "LCC", "LFW"),
           NULL
    )
  }

  # Identify non-trait columns suitable for grouping
  getNonTraitColumns <- function() {
    data <- getActiveInputData()
    # requires available data, not null, to use function
    req(data)
    # vector of traits to be excluded from non-trait cols
    universalTraits <- c(
      "CH", "LDMC", "ldmc", "FS", "FP", "LS", "LDW", "LFW",
      "SLA", "sla", "LA", "PN", "RD", "LNC", "LCC", "succulenceIndex"
    )
    speciesCol <- "species"

    colNames <- colnames(data)
    # identifying and selecting columns from the data that are in (%in%) universalTraits,
    # after converting traits and all cols to lowercase to be case insensitive
    traitColsPresent <- colNames[tolower(colNames) %in% tolower(universalTraits)]
    # setdiff to find differences between columns in the data and the columns from traitcols and species
    nonTraitCols <- setdiff(colNames, c(traitColsPresent, speciesCol))
    return(nonTraitCols)
  }

  # UI Start:

  # Populate the model selection dropdown when the app loads
  # observe is the shiny reactive to events
  observe({
    # list all functions in package
    packageFunctions <- ls("package:CSRcalculator")
    # updates model selection dropdown for the current session. based on that list
    updateSelectInput(session, "functionSelect", choices = packageFunctions)
  })

  # Main UI Rendering:

  # Change main content panel when a topic is selected
  observeEvent(input$introSection, {
    mainContentState(input$introSection)
  })

  # Render the main content panel based on current state
  output$mainContent <- renderUI({
    content <- mainContentState()

    if (content == "intro") {
      return(tagList(
        h3("Introduction"),
        p(HTML("CSR Calculator is a tool designed for ecologists to calculate and assign life strategies to plants based on
                J.P. Grime’s Competitor–Stress-tolerator–Ruderal (CSR) framework
                (<a href='https://www.nature.com/articles/250026a0' target='_blank'>Grime, 1974</a>).
                This framework characterizes species along three strategic axes: competitive ability (C), stress tolerance (S), and ruderality (R),
                reflecting their adaptations to environmental pressures such as resource availability and disturbance."
        )),
        p("This app supports three published models for CSR strategy assignment:"),
        tags$ul(
          tags$li(HTML("<a href='https://www.jstor.org/stable/3546494' target='_blank'>Hodgson et al. (1999)</a>")),
          tags$li(HTML("<a href='https://besjournals.onlinelibrary.wiley.com/doi/full/10.1111/1365-2435.12722' target='_blank'>StrateFy (Pierce et al., 2017)</a>")),
          tags$li(HTML("<a href='https://onlinelibrary.wiley.com/doi/10.1155/2016/1323614' target='_blank'>Morpho-Physiological model (Novakovskiy et al., 2016)</a>"))
        ),

        p("Each model differs in its input requirements, assumptions, and interpretative strengths. You can explore each model in more detail using the Model Guides section in the sidebar."),

        h3("Instructions"),
        p("To get started:"),
        tags$ol(
          tags$li("Upload a dataset containing species and trait measurements for analysis. We currently support .csv, .tsv, .txt and .xlsx filetypes."),
          tags$li("Once uploaded, the app will automatically detect compatible models based on available traits. The app will also detect common variants of expected column names, but you can manually assign columns if needed."),
          tags$li("Use the CSR Analysis panel to choose a model, adjust optional parameters, and process your data."),
          tags$li("View the results in the form of data tables and an interactive ternary plot."),
          tags$li("The Output panel includes optional tools for customizing plots and tables, as well as generating a summary table.")
        ),
        p("Model-specific guidance is provided under Model Guides, as well as an option to load in an example dataset compatible with all three models."),

        h3("Acknowledgments"),
        p("This app was developed as part of a NERC-funded PhD studentship under the ARIES Doctoral Training Partnership As well as in partnership with the Beth Chatto's Gardens"),
        p("For any queries, contact:"),
      ))
    }

    if (content == "strateFy") {
      return(tagList(
        h3("StrateFy Model Guide"),

        p(HTML("The <a href='https://besjournals.onlinelibrary.wiley.com/doi/full/10.1111/1365-2435.12722' target='_blank'>StrateFy model</a>
                (Pierce et al., 2017) was developed using leaf trait data from over 3000 species across 14 different biomes.
                It provides a globally calibrated method for assigning CSR strategies using just three accessible leaf traits:
                specific leaf area (SLA), leaf dry matter content (LDMC), and leaf area (LA). This minimal trait set, coupled with its calibration,
                makes the model especially suitable for analysing large, diverse datasets."
        )),

        p("The model is based on principal component analysis (PCA) of the transformed leaf traits (see table).
           The first PCA axis captures the gradient from competitive to stress-tolerant strategies, driven by variation in
           specific leaf area (SLA) and leaf dry matter content (LDMC). Competitive species tend to have high SLA and low LDMC,
           reflecting investment in rapid growth, while stress-tolerant species show the opposite pattern, with low SLA and
           high LDMC indicating tougher, longer-lived leaves. The second axis, shaped primarily by leaf area, helps distinguish ruderal species,
           which typically have small, short-lived leaves and minimal structural investment."),

        p("Species are positioned along the two PCA axes using regression equations from the global calibration.
           These scores are then converted to C, S, and R percentages, placing each species within the CSR triangle
           and assigning it to one of 19 strategy classes based on the nearest reference point."),

        tags$h4("Required Traits"),
        tableOutput("strateFyTraitTable"),

        p(
          "If SLA is missing, it can be calculated from LA and LDW.",
          br(),
          "If LDMC is missing, it can be calculated from LFW and LDW.",
          br(),
          "We recommend using the LDMC calculations for StrateFy as the model includes an in-built adjustment for succulent species based on the succulence index of each species."
        ),

        div(class = "mt-3",
            p("You can test the StrateFy model with an example dataset:"),
            actionButton("loadStrateFyExample", "Load Example Dataset", class = "btn btn-outline-primary")
        )
      ))
    }

    if (content == "hodgson") {
      return(tagList(
        h3("Hodgson Model Guide"),

        p(HTML("The <a href='https://www.jstor.org/stable/3546494' target='_blank'>Hodgson et al. (1999)</a> model offers one of the first formal attempts
                to classify species within the CSR strategy framework using measurable functional traits. It uses benchmarks derived from field surveys and
                experimental data of British flora, referred to as “gold standards\", to represent idealized examples of the three primary strategies."
        )),
        tags$ul(
          tags$li("The Competitor (C) axis is calibrated using a dominance index, which reflects a species’
                   relative abundance in productive, undisturbed environments. This index reflects the ability of that species to dominate and pre-emptively obtain resources."),

          tags$li("The Stress-tolerator (S) axis is based on principal component analysis (PCA) of growth rates, leaf traits, and nutrient content
                   of plant species grown in low-nutrient environments, which revealed a strong alignment with classic S-strategy traits, such as
                   high LDMC, low SLA, and low nutrient levels. As such, species can be scored along this PCA axis to determine their closeness to an S-type species."),

          tags$li("The Ruderal (R) axis is based on the frequency of ephemeral and monocarpic species found in disturbed sites."),
        ),

        p("To classify a species, the Hodgson model uses up to seven functional traits as predictor variables (see table).
           These traits are transformed and used in regression equations to score each species along the C, S, and R axes.
           The equations will differ between two versions, a non-grasses version including flowering start as an additional variable, and grasses version.
           The resulting scores are converted to coordinates within a CSR triangular space, then matched to the nearest of 19 predefined CSR strategy positions
           based on the smallest sum of squared differences between the species and reference point coordinates."),

        tags$h4("Required Traits"),
        tableOutput("hodgsonTraitTable"),
        p(
          "If SLA is missing, it can be calculated from LA and LDW.",
          br(),
          "If LDMC is missing, it can be calculated from LFW and LDW."
        ),

        div(class = "mt-3",
            p("You can test the Hodgson model with an example dataset:"),
            actionButton("loadHodgsonExample", "Load Example Dataset", class = "btn btn-outline-primary")
        )
      ))
    }

    if (content == "morphoPhys") {
      return(tagList(
        h3("Morpho-Physiological Model Guide"),

        p(HTML("The <a href='https://onlinelibrary.wiley.com/doi/10.1155/2016/1323614' target='_blank'>Morpho-Physiological model</a> (Novakovskiy et al., 2016)
                extends CSR strategy assignment by incorporating both morphological and physiological traits (see table).
                This model was developed using data from 74 boreal and tundra species in northeastern Europe. While this calibration may limit its generality,
                the inclusion of physiological traits enables finer discrimination among species with similar morphology but differing ecological behaviours.")),

        p("To calculate CSR scores, the model first transforms each trait and scores species along two axes derived from PCA analysis of the original dataset.
           These axes reflect underlying ecological gradients: PCA1 generally captures competitive and ruderal tendencies, while PCA2 reflects stress tolerance.
           Weighted combinations of the PCA scores are then used to generate C, S, and R coordinates, which position each species within the CSR triangle.
           Final classification is based on a species’ proximity to 19 predefined strategy coordinates within the triangle, reflecting its closest ecological match."),

        tags$h4("Required Traits"),
        tableOutput("morphoPhysTraitTable"),
        p(
          "If LDMC is missing, it can be calculated from LFW and LDW."
        ),

        div(class = "mt-3",
            p("You can test the Morpho-Physiological model with an example dataset:"),
            actionButton("loadMorphoPhysExample", "Load Example Dataset", class = "btn btn-outline-primary")
        )
      ))
    }

    if (content == "analysis") {
      # Require uploaded data before showing analysis content
      req(originalData())

      # If model not yet applied, show the original input data only
      if (!modelApplied()) {
        return(
          layout_column_wrap(
            width = 1,
            renderOriginalTableCard("Input Data", "downloadOriginal", "csrOriginalTable")
          )
        )
      }

      # If a specific card is expanded (original/processed/plot), show it
      card <- expandedCard()

      if (!is.null(card) && card == "original") {
        return(
          layout_column_wrap(
            width = 1,
            renderExpandedCard("Input Data", "downloadOriginal", "csrOriginalTable")
          )
        )
      }

      if (!is.null(card) && card == "processed") {
        return(
          layout_column_wrap(
            width = 1,
            if (isTRUE(input$showSummaryStats)) {
              # Show processed data + summary stats (in tabs)
              renderExpandedNavCard("downloadCSR", "csrProcessedTable", "csrSummaryTable")
            } else {
              # Show processed data table only
              renderExpandedCard("Processed Data", "downloadCSR", "csrProcessedTable")
            }
          )
        )
      }

      if (!is.null(card) && card == "plot") {
        return(
          layout_column_wrap(
            width = 1,
            renderExpandedPlotCard("Ternary Plot (Expanded)", "csrTernaryPlot")
          )
        )
      }

      # Default layout after model is applied (no specific card expanded)
      tagList(
        layout_column_wrap(
          width = 1,
          if (isTRUE(input$showSummaryStats)) {
            # Processed data + summary stats tabs
            renderCsrNavCard("downloadCSR", "csrProcessedTable", "csrSummaryTable", "expandProcessed")
          } else {
            # Just the processed data table
            renderCsrCard("Processed Data", "downloadCSR", "csrProcessedTable", "expandProcessed")
          }
        ),
        layout_column_wrap(
          width = 1,
          renderCompactPlotCard("Ternary Plot", "csrTernaryPlot", "expandPlot")
        ))
      }
    })

  # Keep mainContent UI alive even when hidden
  outputOptions(output, "mainContent", suspendWhenHidden = FALSE)

  # Trait requirment tables:
  output$hodgsonTraitTable <- renderTable({
    df <- data.frame(
      Abbreviation = c("CH", "LDMC", "FP", "LS", "LDW", "SLA", "FS"),
      Trait = c(
        "Canopy Height",
        "Leaf Dry Matter Content",
        "Flowering Period",
        "Lateral Spread",
        "Leaf Dry Weight",
        "Specific Leaf Area",
        "Flowering Start"
      ),
      Units = c(
        "mm",
        "%",
        "months",
        "categorical (1–6)",
        "mg",
        "mm²/mg",
        "ordinal (1 = March or earlier ... 6 = August or later)"
      ),
      Notes = c(
        "log-transformed",
        "square root",
        "none",
        "none",
        "log + 3",
        "square root",
        "optional: enables non-grass model logic"
      ),
      stringsAsFactors = FALSE
    )
    colnames(df)[4] <- "Notes (transformations handled by model)"
    df
  })

  output$strateFyTraitTable <- renderTable({
    df <- data.frame(
      Abbreviation = c("LA", "SLA", "LDMC"),
      Trait = c(
        "Leaf Area",
        "Specific Leaf Area",
        "Leaf Dry Matter Content"
      ),
      Units = c(
        "mm²",
        "mm²/mg",
        "%"
      ),
      Notes = c(
        "square root",
        "log-transformed",
        "logit-transformed"
      ),
      stringsAsFactors = FALSE
    )
    colnames(df)[4] <- "Notes (transformations handled by model)"
    df
  })

  output$morphoPhysTraitTable <- renderTable({
    df <- data.frame(
      Abbreviation = c("CH", "LDMC", "FP", "LS", "LDW", "PN", "RD", "LNC", "LCC"),
      Trait = c(
        "Canopy Height",
        "Leaf Dry Matter Content",
        "Flowering Period",
        "Lateral Spread",
        "Leaf Dry Weight",
        "Net Photosynthesis",
        "Dark Respiration Rate",
        "Leaf Nitrogen Content",
        "Leaf Carbon Concentration"
      ),
      Units = c(
        "mm",
        "%",
        "months",
        "categorical (1–6)",
        "mg",
        "mg CO₂ / g dry weight / hour",
        "mg CO₂ / g dry weight / hour",
        "mg/g",
        "mg/g"
      ),
      Notes = c(
        "log-transformed",
        "square root",
        "none",
        "none",
        "log + 3",
        "square root",
        "square root",
        "square root",
        "square root"
      ),
      stringsAsFactors = FALSE
    )
    colnames(df)[4] <- "Notes (transformations handled by model)"
    df
  })

  # Helper Functions:

  # Load an example dataset and update app state
  loadExampleDataset <- function(filePath, modelName) {
    if (!file.exists(filePath)) {
      showNotification(paste(modelName, "example dataset not found."), type = "error")
      return(NULL)
    }
    # fread for faster csv reading
    data <- data.table::fread(filePath, data.table = FALSE)
    data <- renameColumns(data)

    resetAppState()
    originalData(data)
    session$userData$originalUploadedData <- data
    updateCompatibleModels(data)
    # switch main content state to analysis from intro / selected topic
    mainContentState("analysis")
    updateSelectInput(session, "functionSelect", selected = modelName)

    showNotification(paste("Loaded example dataset for", modelName, "model."), type = "message")
  }

  # Example Dataset Loading:

  # Load example dataset for StrateFy model
  observeEvent(input$loadStrateFyExample, {
    loadExampleDataset("data/exampleData.csv", "strateFy")
  })

  # Load example dataset for Hodgson model
  observeEvent(input$loadHodgsonExample, {
    loadExampleDataset("data/exampleData.csv", "hodgson")
  })

  # Load example dataset for Morpho-Phys model
  observeEvent(input$loadMorphoPhysExample, {
    loadExampleDataset("data/exampleData.csv", "morphoPhys")
  })

  # File Upload Handling:

  # Switch to analysis view after upload
  observeEvent(input$fileInput, {
    data <- readValidateData(input, session)
    if (!is.null(data)) {
      resetAppState()
      originalData(data)
      session$userData$originalUploadedData <- data
      updateCompatibleModels(data)

      mainContentState("analysis")

      # Clear active link from intro sidebar (green highlight)
      session$sendCustomMessage("clearActiveSidebarLink", list())
    }
  })

  # Render list of compatible models
  output$compatibleModelsText <- renderText({
    req(compatibleModels())
    paste("Compatible models:", paste(compatibleModels(), collapse = ", "))
  })

  # Render the uploaded data set
  output$csrOriginalTable <- DT::renderDataTable({
    req(originalData())
    data <- getActiveInputData()

    DT::datatable(
      data,
      # render options
      filter = "top",
      rownames = FALSE,
      selection = "multiple",
      options = list(
        dom = '<"d-flex justify-content-between align-items-center"lfB>rt<"bottom"ip>',
        pageLength = 10,
        lengthMenu = list(c(5, 10, 25, 50, -1), c("5", "10", "25", "50", "All")),
        autoWidth = TRUE,
        responsive = TRUE,
        search = list(regex = TRUE),
        columnDefs = list(
          list(orderSequence = c("desc", "asc"), targets = "_all"),
          list(className = "dt-center", targets = "_all")
        )
      )
    ) %>%
      DT::formatRound(
        columns = names(data)[sapply(data, is.numeric)],
        digits = 2
      )
  })

  # Manual Column Assignment:

  # Render UI for manual column selection
  output$columnSelectionUI <- renderUI({
    # check data is available before proceeding
    req(originalData(), input$manualColumnSelection)
    # populate dropdown lists for each trait with data column names
    columnChoices <- names(originalData())
    # display traits based on selected model
    selectedModel <- input$manualTraitModel %||% input$functionSelect
    traits <- getRequiredTraits(selectedModel)
    # fallback
    if (is.null(traits)) {
      return(p("Please select a model to assign traits."))
    }

    # Split traits into two per row instead of one long column
    rows <- split(traits, ceiling(seq_along(traits) / 2))

    tagList(
      selectInput("manualTraitModel", "Model for Trait Assignment",
                  choices = c("hodgson", "strateFy", "morphoPhys"),
                  selected = selectedModel
      ),
      lapply(rows, function(traitPair) {
        fluidRow(
          lapply(traitPair, function(trait) {
            column(
              width = 6,
              selectInput(
                # create input IDs based on user input, e.g. user selects LA, ID = trait_LA, paste0 combines "trait)_" and selected column or trait
                inputId = paste0("trait_", trait),
                label = trait,
                choices = c("", columnChoices),
                selected = ""
              )
            )
          })
        )
      }),
      div(class = "mt-2 d-grid gap-0",
          actionButton("applyColumnChanges", "Apply Trait Assignment", class = "btn btn-outline-dark mb-1"),
          actionButton("resetColumns", "Reset", class = "btn btn-outline-secondary")
      )
    )
  })

  # Apply manual trait assignments
  observeEvent(input$applyColumnChanges, {
    req(originalData(), input$manualTraitModel)
    # define data and traits from trait lists
    data <- originalData()
    traits <- getRequiredTraits(input$manualTraitModel)

    # Only use non-empty trait selections, so user doesn't have to rename for all traits
    # sapply applies function to each element of traits list/vector, paste0... calls the user input ID, creating a vector of traits and input IDs
    validAssignments <- sapply(traits, function(trait) input[[paste0("trait_", trait)]])
    # valid assignments are not equal to nothing, allows the user to keep traits empty
    validAssignments <- validAssignments[validAssignments != ""]

    # Check for duplicate trait assignments
    if (length(validAssignments) != length(unique(validAssignments))) {
      showNotification("Each trait must be assigned to a unique column.", type = "error")
      return(NULL)
    }

    # Temporarily rename selected columns
    # loop through each element in the assignment vector
    for (trait in names(validAssignments)) {
      # take the selected column for renaming
      col <- validAssignments[[trait]]
      # check the column they selected exists in their data
      if (col %in% names(data)) {
        # temporary renmaming by adding temp as a prefix to the selected column
        names(data)[names(data) == col] <- paste0(".__temp__", trait)
      }
    }

    # Assign standard trait names
    # again looping through each element
    for (trait in names(validAssignments)) {
      # look for temp prefix and replace column name with the standard version
      names(data)[names(data) == paste0(".__temp__", trait)] <- trait
    }

    renamedData(data)
    manualSelectionActive(TRUE)
    updateCompatibleModels(data)
    showNotification("Trait assignments applied.", type = "message")
  })

  # Reset trait assignments back to original
  observeEvent(input$resetColumns, {
    req(originalData())
    renamedData(originalData())
    manualSelectionActive(FALSE)
    updateCompatibleModels(originalData())

    # Clear downstream transformations
    traitAveragedData(NULL)
    processedModelData(NULL)
    csrAveragedData(NULL)
    modelApplied(FALSE)
  })

  # Trait Averaging:

  # Render dropdown for group by column (for trait averaging)
  output$traitGroupByUI <- renderUI({
    req(originalData())

    data <- getActiveInputData()
    # grouping options should not be trait columns
    groupCandidates <- getNonTraitColumns()

    # Allow character, factor, or numeric columns with < 20 unique values
    validGroups <- groupCandidates[
      sapply(data[groupCandidates], function(col) {
        is.character(col) ||
          is.factor(col) ||
          (is.numeric(col) && length(unique(na.omit(col))) <= 20)
      })
    ]

    if (length(validGroups) == 0) {
      return(tags$em("No suitable grouping columns available."))
    }

    selectInput(
      inputId = "groupByColumn",
      label = "Group Traits By:",
      choices = validGroups,
      selected = validGroups[1]
    )
  })

  # Render trait selection UI
  output$traitSelectionUI <- renderUI({
    req(originalData(), input$averagingMode == "traits")
    data <- getActiveInputData()
    numericCols <- names(data)[sapply(data, is.numeric)]

    tagList(
      uiOutput("traitGroupByUI"),
      selectInput("traitsToAverage", "Select Traits to Average", choices = numericCols, multiple = TRUE),
      actionButton("applyAveraging", "Apply Averages"),
      actionButton("resetAveraging", "Reset")
    )
  })

  # Apply trait averaging based on selected group and traits
  observeEvent(input$applyAveraging, {
    req(originalData(), input$traitsToAverage, input$groupByColumn)
    data <- getActiveInputData()
    groupByCol <- input$groupByColumn

    # Safety checks
    if (!(groupByCol %in% names(data))) {
      showNotification(paste("Error: Group By column not found:", groupByCol), type = "error")
      return(NULL)
    }

    if (length(input$traitsToAverage) == 0) {
      showNotification("Error: No traits selected for averaging.", type = "error")
      return(NULL)
    }

    # Average the selected traits
    averaged <- data %>%
      # dplyr function to group data frame by the whole selected column
      group_by(.data[[groupByCol]]) %>%
      # dplyr summarise function summarises data into one row per group, "across" all of the traits user selected to average, only using mean
      summarise(across(all_of(input$traitsToAverage), ~ mean(.x, na.rm = TRUE)), .groups = "drop")

    # Add back reserved columns (first value from each group), for columns not selected for averaging
    reservedCols <- setdiff(names(data), c(input$traitsToAverage, groupByCol))
    metadata <- data %>%
      group_by(.data[[groupByCol]]) %>%
      summarise(across(all_of(reservedCols), ~ first(.x)), .groups = "drop")

    finalAveraged <- left_join(averaged, metadata, by = groupByCol)

    traitAveragedData(finalAveraged)
    averagingActive(TRUE)
    updateCompatibleModels(finalAveraged)
  })

  # Reset trait averaging
  observeEvent(input$resetAveraging, {
    originalData(session$userData$originalUploadedData)
    traitAveragedData(NULL)
    averagingActive(FALSE)
    processedModelData(NULL)
    csrAveragedData(NULL)
    modelApplied(FALSE)
    updateCompatibleModels(originalData())
  })

  # CSR Averaging (Scores/Percentages):

  # Helper to get non-trait columns for grouping
  getNonTraitColumns <- function() {
    data <- getActiveInputData()
    req(data)

    # Universal list of trait columns across models
    universalTraits <- c(
      "CH", "LDMC", "ldmc", "FS", "FP", "LS", "LDW", "LFW",
      "SLA", "sla", "LA", "PN", "RD", "LNC", "LCC", "succulenceIndex"
    )
    speciesCol <- "species"

    colNames <- colnames(data)
    traitColsPresent <- colNames[tolower(colNames) %in% tolower(universalTraits)]
    nonTraitCols <- setdiff(colNames, c(traitColsPresent, speciesCol))
    return(nonTraitCols)
  }

  # Render UI for selecting a grouping column for CSR score averaging
  output$csrGroupByUI <- renderUI({
    req(originalData())

    data <- getActiveInputData()
    groupCandidates <- getNonTraitColumns()

    # Allow character, factor, or numeric columns with fewer than 20 unique values
    validGroups <- groupCandidates[
      sapply(data[groupCandidates], function(col) {
        is.character(col) ||
          is.factor(col) ||
          (is.numeric(col) && length(unique(na.omit(col))) <= 20)
      })
    ]

    if (length(validGroups) == 0) {
      return(tags$em("No suitable grouping columns available."))
    }

    selectInput(
      inputId = "csrGroupBy",
      label = "Group CSR scores by:",
      choices = validGroups,
      selected = validGroups[1]
    )
  })

  # Display CSR averaging status message
  output$csrAveragingStatus <- renderText({
    req(csrAveragingConfirmed())
    group <- input$csrGroupBy
    if (nzchar(group)) {
      paste("Averaging CSR scores by:", group, "\n(Will apply after model is run)")
    } else {
      "No grouping column set for CSR averaging."
    }
  })

  # Confirm CSR score averaging
  observeEvent(input$applyCsrAveraging, {
    req(originalData(), input$csrGroupBy)
    data <- getActiveInputData()
    groupCol <- input$csrGroupBy

    # Safety checks before confirming CSR averaging
    if (!(groupCol %in% names(data))) {
      showNotification("Error: Group column not found.", type = "error")
      return(NULL)
    }

    if (all(is.na(data[[groupCol]]))) {
      showNotification("Error: Group column is entirely NA.", type = "error")
      return(NULL)
    }

    if (length(unique(na.omit(data[[groupCol]]))) <= 1) {
      showNotification("Error: Group column does not have multiple groups.", type = "error")
      return(NULL)
    }

    if (!(is.character(data[[groupCol]]) || is.factor(data[[groupCol]]) || is.numeric(data[[groupCol]]))) {
      showNotification("Error: Group column must be character, factor, or numeric.", type = "error")
      return(NULL)
    }

    # Passed all checks, allow CSR averaging
    csrAveragingConfirmed(TRUE)
  })

  # Reset CSR averaging
  observeEvent(input$resetCsrAveraging, {
    csrAveragingConfirmed(FALSE)
  })


  # Reset logic if user changes averaging mode
  observeEvent(input$averagingMode, {
    originalData(session$userData$originalUploadedData)

    if (input$averagingMode == "csr") {
      # Switching to CSR score averaging
      traitAveragedData(NULL)
      modelApplied(FALSE)
      processedModelData(NULL)
      csrAveragedData(NULL)
      averagingActive(FALSE)

    } else if (input$averagingMode == "traits") {
      # Switching to trait averaging
      csrAveragedData(NULL)
      modelApplied(FALSE)
      processedModelData(NULL)

    } else if (input$averagingMode == "none") {
      # No averaging
      traitAveragedData(NULL)
      csrAveragedData(NULL)
      processedModelData(NULL)
      modelApplied(FALSE)
      averagingActive(FALSE)
    }
  })

  # Apply selected CSR model to the active input data
  observeEvent(input$applyModel, {
    req(input$functionSelect)
    data <- getActiveInputData(); req(data)

    # Validate that all required inputs are available
    validationError <- validateCsrModel(data, input$functionSelect, input)
    if (!is.null(validationError)) {
      showNotification(validationError, type = "error")
      return(NULL)
    }

    data <- removeDuplicateColumns(data)  # Clean data before model

    # Run the selected model
    result <- NULL
    if (input$functionSelect == "strateFy") {
      # strateFy intermediate trait calc options, default is true, calculate them
      calcSLA  <- !isTRUE(input$useProvidedSLA)
      calcLDMC <- !isTRUE(input$useProvidedLDMC)
      # remove these columns if the user chose to let the app calculate them
      if (calcSLA)  data <- dplyr::select(data, -any_of("SLA"))
      if (calcLDMC) data <- dplyr::select(data, -any_of(c("LDMC", "succulenceIndex")))
      result <- tryCatch(
        strateFy(data, calcSLA = calcSLA, calcLDMC = calcLDMC),
        error = function(e) paste("Error applying StrateFy model:", e$message)
      )
    } else if (input$functionSelect == "hodgson") {
      # intermediate trait calc option, default is false, do not calc
      calcSLA  <- isTRUE(input$hodgsonCalculateSLA)
      calcLDMC <- isTRUE(input$hodgsonCalculateLDMC)
      # remove these columns if the user chose to let the app calculate them
      if (calcSLA)  data <- dplyr::select(data, -any_of("SLA"))
      if (calcLDMC) data <- dplyr::select(data, -any_of("LDMC"))
      # grass or non-grass where available option
      preferNonGrasses <- isTRUE(input$hodgsonPreferNonGrasses)
      result <- tryCatch(
        hodgson(data, calcSLA = calcSLA, calcLDMC = calcLDMC, preferNonGrasses = preferNonGrasses),
        error = function(e) paste("Error applying Hodgson model:", e$message)
      )
    } else if (input$functionSelect == "morphoPhys") {
      # intermediate trait calc option, default is false, do not calc
      calcLDMC <- isTRUE(input$morphoPhysCalculateLDMC)
      # remove these columns if the user chose to let the app calculate them
      if (calcLDMC) data <- dplyr::select(data, -any_of("LDMC"))
      result <- tryCatch(
        morphoPhys(data, calcLDMC = calcLDMC),
        error = function(e) paste("Error applying Morpho-Phys model:", e$message)
      )
    } else {
      result <- applyCsrModel(data, input$functionSelect, input)
    }

    # Show error message if model failed
    if (is.character(result)) {
      showNotification(result, type = "error")
      return(NULL)
    }

    # Store processed results
    result <- removeDuplicateColumns(result)
    processedModelData(result)

    # Optionally apply CSR averaging
    if (input$averagingMode == "csr" && csrAveragingConfirmed()) {
      groupCols <- unlist(strsplit(input$csrGroupBy, ",\\s*"))
      averaged <- switch(
        input$functionSelect,
        "hodgson"    = averageHodgsonCsrByGroup(result, groupCols),
        "morphoPhys" = averageMorphoPhysCsrByGroup(result, groupCols),
        "strateFy"   = averageStrateFyCsrByGroup(result, groupCols)
      )
      if (is.null(averaged)) {
        showNotification("Error: model not supported for CSR score averaging.", type = "error")
        return(NULL)
      }
      csrAveragedData(removeDuplicateColumns(averaged))
    } else {
      csrAveragedData(result)
    }

    # Flag that model output is ready
    modelApplied(TRUE)
  })

  # Render the processed model output (or CSR-averaged output) table
  output$csrProcessedTable <- DT::renderDataTable({
    data <- if (!is.null(csrAveragedData())) csrAveragedData() else processedModelData()
    req(data)

    data <- removeDuplicateColumns(data)

    # Reorder columns: species -> non-trait columns -> trait columns
    traitCols <- c("CH", "LDMC", "ldmc", "FS", "FP", "LS", "LDW", "LFW", "SLA", "sla", "LA", "PN", "RD", "LNC", "LCC", "succulenceIndex")
    colNames <- colnames(data)
    traitColsPresent <- colNames[tolower(colNames) %in% tolower(traitCols)]
    speciesCol <- if ("species" %in% colNames) "species" else NULL
    nonTraitCols <- setdiff(colNames, c(traitColsPresent, speciesCol))
    reorderedCols <- c(speciesCol, nonTraitCols, traitColsPresent)
    reorderedCols <- reorderedCols[!is.na(reorderedCols)]

    data <- data[, reorderedCols, drop = FALSE]

    # Optionally hide trait columns
    if (isTruthy(input$hideTraitColumns)) {
      data <- data[, !colnames(data) %in% traitColsPresent, drop = FALSE]
    }

    DT::datatable(
      data,
      filter = "top",
      rownames = FALSE,
      selection = "multiple",
      options = list(
        dom = '<"d-flex justify-content-between align-items-center"lfB>rt<"bottom"ip>',
        pageLength = tablePageLength(),
        lengthMenu = list(c(5, 10, 25, 50, -1), c("5", "10", "25", "50", "All")),
        autoWidth = TRUE,
        search = list(regex = TRUE),
        columnDefs = list(
          list(orderSequence = c("desc", "asc"), targets = "_all"),
          list(className = "dt-center", targets = "_all")
        )
      )
    ) %>%
      DT::formatRound(columns = names(data)[sapply(data, is.numeric)], digits = 2)
  })

  # Update table page length reactively
  observeEvent(input$csrProcessedTable_length, {
    tablePageLength(input$csrProcessedTable_length)
  })

  # Render the summary statistics table
  output$csrSummaryTable <- DT::renderDT({
    req(processedModelData())

    data <- if (!is.null(csrAveragedData())) csrAveragedData() else processedModelData()
    req(data)

    groupCol <- input$summaryGroupBy
    if (is.null(groupCol) || groupCol == "None") groupCol <- NULL

    statsDF <- buildSummaryStats(data, groupBy = groupCol)
    req(statsDF)

    numericCols <- names(statsDF)[sapply(statsDF, is.numeric)]

    DT::datatable(
      statsDF,
      rownames = FALSE,
      selection = "none",
      options = list(
        dom = "t",
        pageLength = nrow(statsDF)
      )
    ) %>%
      DT::formatRound(columns = numericCols, digits = 2)
  })

  # Render UI for selecting grouping variable for summary statistics
  output$summaryGroupByUI <- renderUI({
    req(input$showSummaryStats)

    data <- if (!is.null(csrAveragedData())) csrAveragedData() else processedModelData()
    req(data)

    # Grouping candidates: character or factor columns (except strategyClass)
    groupCandidates <- names(data)[sapply(data, function(col) is.character(col) || is.factor(col))]
    groupCandidates <- setdiff(groupCandidates, "strategyClass")

    if (length(groupCandidates) == 0) {
      return(tags$em("No suitable grouping columns available."))
    }

    selectInput(
      inputId = "summaryGroupBy",
      label = "Group summary by:",
      choices = c("None", groupCandidates),
      selected = "None"
    )
  })

  # Render ternary plot
  output$csrTernaryPlot <- renderPlotly({
    req(redrawPlot())

    # Use processed or CSR-averaged data
    fullData <- if (!is.null(csrAveragedData())) {
      csrAveragedData()
    } else {
      processedModelData()
    }
    req(fullData)

    # Filter data by visible rows in the table if option is enabled
    if (isTRUE(input$filterPlotByTable)) {
      rowIdx <- input$csrProcessedTable_rows_all
      if (is.null(rowIdx)) rowIdx <- integer(0)
      filteredData <- fullData[rowIdx, , drop = FALSE]

      highlightRows <- rep(FALSE, nrow(filteredData))
      if (!is.null(input$csrProcessedTable_rows_selected)) {
        highlightRows[input$csrProcessedTable_rows_selected] <- TRUE
      }
    } else {
      filteredData <- fullData

      highlightRows <- rep(FALSE, nrow(filteredData))
      if (!is.null(input$csrProcessedTable_rows_selected)) {
        highlightRows[input$csrProcessedTable_rows_selected] <- TRUE
      }
    }

    # Generate plot object
    generateCsrPlotObject(filteredData, input, highlightRows)
  })

  # Update available columns for shape mapping
  observe({
    req(csrAveragedData() %||% processedModelData())
    data <- csrAveragedData() %||% processedModelData()

    possibleShapeCols <- names(data)[sapply(names(data), function(colname) {
      colData <- data[[colname]]
      (is.character(colData) || is.factor(colData) || is.numeric(colData)) &&
        length(unique(na.omit(colData))) <= 5
    })]

    updateSelectInput(
      session,
      "shapeBy",
      choices = c("None", possibleShapeCols),
      selected = "None"
    )
  })

  # Show/hide UI elements based on app state
  output$csrShowOriginalTable <- reactive({ !is.null(originalData()) })
  output$csrShowProcessedElements <- reactive({
    modelApplied() && (!is.null(csrAveragedData()) || !is.null(processedModelData()))
  })
  outputOptions(output, "csrShowOriginalTable", suspendWhenHidden = FALSE)
  outputOptions(output, "csrShowProcessedElements", suspendWhenHidden = FALSE)

  # Handle card expansion and collapse events
  observeEvent(input$expandOriginal, {
    expandedCard("original")
  })

  observeEvent(input$expandProcessed, {
    expandedCard("processed")
  })

  observeEvent(input$expandPlot, {
    expandedCard("plot")
    redrawPlot(redrawPlot() + 1)
    session$sendCustomMessage("forceRelayout", list(id = "csrTernaryPlot"))
  })

  observeEvent(input$collapse, {
    expandedCard(NULL)
    redrawPlot(redrawPlot() + 1)
    session$sendCustomMessage("forceRelayout", list(id = "csrTernaryPlot"))
  })

  # Download original data
  output$downloadOriginal <- downloadHandler(
    filename = function() {
      suffix <- if (!is.null(traitAveragedData())) {
        "_averaged"
      } else if (manualSelectionActive()) {
        "_renamed"
      } else {
        ""
      }
      paste0("Original_CSR_Table", suffix, "_", Sys.Date(), ".csv")
    },
    content = function(file) {
      data <- getActiveInputData()
      write.csv(data, file, row.names = FALSE)
    }
  )

  # Download processed or summary data
  output$downloadCSR <- downloadHandler(
    filename = function() {
      tabName <- input$csrTabs %||% input$csrTabsExpanded %||% "Processed"
      prefix <- if (tabName == "Summary") "Summary_CSR_Table" else "Processed_CSR_Table"
      paste0(prefix, "_", Sys.Date(), ".csv")
    },
    content = function(file) {
      tabName <- input$csrTabs %||% input$csrTabsExpanded %||% "Processed"

      if (tabName == "Summary") {
        groupCol <- input$summaryGroupBy
        if (is.null(groupCol) || groupCol == "None") groupCol <- NULL

        data <- if (!is.null(csrAveragedData())) {
          csrAveragedData()
        } else {
          processedModelData()
        }
        req(data)
        data <- removeDuplicateColumns(data)
        summaryData <- buildSummaryStats(data, groupBy = groupCol)
        write.csv(summaryData, file, row.names = FALSE)
      } else {
        data <- if (!is.null(csrAveragedData())) {
          csrAveragedData()
        } else {
          processedModelData()
        }
        req(data)
        write.csv(data, file, row.names = FALSE)
      }
    }
  )
}
