## Supplementary material for "CSR Calculator: An R package and Shiny application for assigning plant ecological strategies using trait data": R code text files used in the CSRcalculator R package and shiny application.: hodgson.txt

#' Calculate CSR Scores and Assign a CSR Strategy Using the Hodgson et al. (1999) Model
#'
#' This function calculates and assigns CSR strategy types to plant species
#' based on the Hodgson et al. (1999) model, as well as performing
#' the required trait transformations for the model.
#' It implements two versions of the model (for grasses and non-grasses) and can automatically select the appropriate equations based on the presence of valid flowering start (FS) values.
#'
#' @param data A data frame containing species names and trait data.
#' FS determines whether the grass or non-grass version of the model is used.
#' LFW and LA may be provided instead of LDMC and SLA, respectively.
#' \describe{
#'   \item{CH}{Canopy height (mm)}
#'   \item{LDMC}{Leaf dry matter content (\%)}
#'   \item{FP}{Flowering period (months)}
#'   \item{LS}{Lateral spread (categorical scale: 1–6, based on Hodgson et al. 1999)}
#'   \item{LDW}{Leaf dry weight (mg)}
#'   \item{SLA}{Specific leaf area (mm²/mg)}
#'   \item{LFW}{Leaf fresh weight (mg)}
#'   \item{LA}{Leaf area (mm²)}
#'   \item{FS}{Flowering start(1-6, 1 = before or during March, 2 = April ... 6 = August or after)}
#' }
#' @param calcLDMC Logical. Whether to calculate LDMC from LDW and LFW. Default is FALSE.
#' @param calcSLA Logical. Whether to calculate SLA from LA and LDW. Default is FALSE.
#' @param preferNonGrasses Logical. If TRUE, the function uses the non-grass equations wherever valid FS values (1–6) are present. If FALSE (default), the function uses only the grass equations.
#'
#' @return A data frame with CSR scores (-2.5 to 2.5), CSR percentages (C%, S%, R%), and assigned strategy classification.
#'
#' @import dplyr
#' @export
#'
#' @examples
#' # hodgson(exampleData)

hodgson <- function(data, calcLDMC = FALSE, calcSLA = FALSE, preferNonGrasses = FALSE) {
  # column renaming helper function
  renameColumnsHodgson <- function(data) {
    # making a list for names = required columns, values are regex patterns that might appear in a data set
    # list allows for multiple regexes
    renameDict <- list(
      # e.g. 'species' is the target name, telling the function that the values are equivalent
      # ^ and $ get exact matches so any word with sp in it will not match, '[._ ]?' matches users choice of spacing
      "species" = c("^species$", "^sp\\.$", "^sp$", "^taxon$", "^species[._ ]?name$", "^taxon[._ ]?name$"),
      "CH" = c("^canopy[._ ]?height$", "^CH$"),
      "LDMC" = c("^leaf[._ ]?dry[._ ]?matter[._ ]?content$", "^LDMC$"),
      "FP" = c("^flowering[._ ]?period$", "^FP$"),
      "LS" = c("^lateral[._ ]?spread$", "^LS$"),
      "LDW" = c("^leaf[._ ]?dry[._ ]?weight$", "^LDW$"),
      "LFW" = c("^leaf[._ ]?fresh[._ ]?weight$", "^LFW$"),
      "SLA" = c("^specific[._ ]?leaf[._ ]?area$", "^SLA$"),
      "LA" = c("^leaf[._ ]?area$", "^LA$"),
      "FS" = c("^flowering[._ ]?start$", "^FS$")
    )
    # for each user column loop through each name in dictionary (renameDict)
    for (colName in names(renameDict)) {
      # use grep() to find and return user columns matching a single regex patterns
      # patterns are taken from renameDict[[colName]] and collapsed into one regex string with paste(..., collapse = "|") to work with grep
      matchedCols <- grep(paste(renameDict[[colName]], collapse = "|"), names(data), ignore.case = TRUE, value = TRUE)
      # if at least one column matches...
      if (length(matchedCols) > 0) {
        # rename the matching column to the name in the dictionary list
        names(data)[names(data) %in% matchedCols] <- colName
      }
    }

    return(data)
  }
  # rename columns
  data <- renameColumnsHodgson(data)

  # check for species column
  if (!"species" %in% names(data)) {
    stop("data must contain a 'species' column")
  }

  # determine required traits (not checking for FS yet, so it is optional)
  if (calcLDMC && calcSLA) {
    requiredCols <- c("species", "CH", "FP", "LS", "LDW", "LFW", "LA")
  } else if (calcLDMC) {
    requiredCols <- c("species", "CH", "FP", "LS", "LDW", "LFW", "SLA")
  } else if (calcSLA) {
    requiredCols <- c("species", "CH", "FP", "LS", "LDW", "LDMC", "LA")
  } else {
    requiredCols <- c("species", "CH", "FP", "LS", "LDW", "LDMC", "SLA")
  }

  # remove rows with missing values
  # FS again not included as not all rows may have a value (i.e. mix of grasses and non-grasses)
  # pipeline to filter user data through - if any required columns contain na
  missingRows <- data %>% filter(if_any(all_of(requiredCols), ~ is.na(.)))
  if (nrow(missingRows) > 0) {
    # paste species names of rows with missing values into a vector then collapse into a string to be readable
    # e.g. (from c("species1",... to species1,...))
    removedSpecies <- paste(unique(missingRows$species), collapse = ", ")
    # display warning
    warning(sprintf(
      "%d row(s) removed due to missing trait values. affected species: %s",
      nrow(missingRows),
      removedSpecies
    ))
  }
  # anti_join to keep rows that do not match defined missingrows based on species column
  data <- anti_join(data, missingRows, by = "species")

  # optional LDMC and SLA calculations
  # adding columns via mutate
  if (calcLDMC) data <- data %>% mutate(LDMC = (LDW * 100) / LFW)
  if (calcSLA)  data <- data %>% mutate(SLA = LA / LDW)

  # model selection logic
  # if FS is a column, convert to characters (as.character) then numeric (as.numeric) whilst ignoring warnings
  # standardizes column to numbers or NA, else create a vector column of NA matching number of rows to rest of data so FS can be flagged as missing to use grasses version instead
  fsNumeric <- if ("FS" %in% names(data)) suppressWarnings(as.numeric(as.character(data$FS))) else rep(NA_real_, nrow(data))
  # if preferNonGrasses is True, check non na (!is.na) FS is numeric and between 1 and 6
  isNonGrass <- if (preferNonGrasses) (!is.na(fsNumeric) & fsNumeric %in% 1:6) else rep(FALSE, nrow(data))

  # trait transformations
  transformed <- data %>%
    mutate(
      chPr  = case_when(
        CH > 999 ~ 6,
        CH > 599 ~ 5,
        CH > 299 ~ 4,
        CH > 99  ~ 3,
        CH > 49  ~ 2,
        TRUE     ~ 1
      ),
      ldmcPr = sqrt(LDMC),
      fpPr   = FP,
      fsPr   = suppressWarnings(as.numeric(as.character(FS))),
      lsPr   = LS,
      ldwPr  = log(LDW) + 3,
      slaPr  = sqrt(SLA)
    )

  # create raw scores columns
  transformed$rawC <- NA_real_
  transformed$rawS <- NA_real_
  transformed$rawR <- NA_real_

  # grass model
  # where isNonGrass = False, select those rows and the placeholder columns, assign them to the pipeline
  transformed[!isNonGrass, c("rawC", "rawS", "rawR")] <- transformed[!isNonGrass, ] %>%
    # mutate placeholder columns and the selected rows to contain raw scores for grasses
    mutate(
      rawC = (0.141 * chPr^2) + (0.09061 * lsPr^2),
      rawS = 54.6 - (1.666 * chPr^2) + (1.069 * ldmcPr^2) - (2.732 * slaPr^2) + (1.722 * lsPr^2),
      rawR = (2.518 * fpPr) - (2.748 * ldwPr) + (5.37 * slaPr)
      # select score columns only, remove others
    ) %>% dplyr::select(rawC, rawS, rawR)

  # non-grass model
  transformed[isNonGrass, c("rawC", "rawS", "rawR")] <- transformed[isNonGrass, ] %>%
    mutate(
      rawC = (0.09245 * chPr^2) + (0.05631 * lsPr^2) + (0.01595 * ldwPr^2),
      rawS = -39.52 - (7.581 * chPr) + (2.633 * ldmcPr^2) - (0.351 * ldwPr^2),
      rawR = -(1.158 * ldmcPr^2) + (3.137 * fpPr) + (3.145 * fsPr) -
        (0.0849 * ldwPr^2) - (1.193 * slaPr^2) + (11.4 * slaPr)
    ) %>% dplyr::select(rawC, rawS, rawR)

  # score transformations and capping
  transformed <- transformed %>%
    mutate(
      cScoreRaw = -2.5 + 0.839 * rawC,
      sScoreRaw = if_else(isNonGrass, -1.249 + 0.0531 * rawS, -1.103 + 0.0474 * rawS),
      rScoreRaw = -2.5 + 0.119 * rawR,
      # clip raw scores between -2.5 and 2.5 then round to nearest 10th
      cScore = trunc(pmin(pmax(cScoreRaw, -2.5), 2.5) * 10) / 10,
      sScore = trunc(pmin(pmax(sScoreRaw, -2.5), 2.5) * 10) / 10,
      rScore = trunc(pmin(pmax(rScoreRaw, -2.5), 2.5) * 10) / 10
    )

  # convert to percentages
  transformed <- transformed %>%
    mutate(
      cPercent = 100 * (cScore + 2.5) / (cScore + sScore + rScore + 7.5),
      sPercent = 100 * (sScore + 2.5) / (cScore + sScore + rScore + 7.5),
      rPercent = 100 * (rScore + 2.5) / (cScore + sScore + rScore + 7.5)
    )

  # strategy classification
  # create a data frame to assign CSR value Strategy classes
  ref <- data.frame(
    strategy = c("C", "C/CR", "C/SC", "CR", "C/CSR", "SC", "CR/CSR", "SC/CSR", "R/CR",
                 "CSR", "S/SC", "R/CSR", "S/CSR", "R", "SR/CSR", "S", "R/SR", "S/SR", "SR"),
    cRef = c(2, 1, 1, 0, 1, 0, 0, 0, -1, 0, -1, -1, -1, -2, -1, -2, -2, -2, -2),
    sRef = c(-2, -2, -1, -2, -1, 0, -1, 0, -2, 0, 1, -1, 1, -2, 0, 2, -1, 1, 0),
    rRef = c(-2, -1, -2, 0, -1, -2, 0, -1, 1, 0, -2, 1, -1, 2, 0, -2, 1, -1, 0)
  )
  # classify function for c s and r scores (in use c = cScore etc.)
  classify <- function(c, s, r) {
    # distance between c s and r scores (coordinates) and reference c s and r from referencde data frame
    d <- sqrt((ref$cRef - c)^2 + (ref$sRef - s)^2 + (ref$rRef - r)^2)
    # which.min finds row number with smallest distance, square brackets to return strategy name out of ref data frame
    ref$strategy[which.min(d)]
  }

  transformed <- transformed %>%
    rowwise() %>%
    # applying classify function to the transformed data scores. add column to data set
    mutate(strategyClass = classify(cScore, sScore, rScore)) %>%
    ungroup()

  # optional model version column
  if ("FS" %in% names(data)) {
    transformed <- transformed %>%
      mutate(hodgsonModelVersion = if_else(isNonGrass, "non-grass", "grass"))
  }

  # select output columns
  outCols <- c("cScore", "sScore", "rScore", "cPercent", "sPercent", "rPercent", "strategyClass")
  if (calcLDMC) outCols <- c("LDMC", outCols)
  if (calcSLA)  outCols <- c("SLA", outCols)
  if ("hodgsonModelVersion" %in% names(transformed)) {
    outCols <- c("hodgsonModelVersion", outCols)
  }

  result <- cbind(data, transformed %>% dplyr::select(dplyr::any_of(outCols)))
  return(result)
}
