## Supplementary material for "CSR Calculator: An R package and Shiny application for assigning plant ecological strategies using trait data": R code text files used in the CSRcalculator R package and shiny application.: morphoPhys.txt

#' Calculate CSR Scores and Assign a CSR Strategy Using the Morphophysiological Model (Novakovskiy et al. 2016)
#'
#' This function calculates and assigns CSR strategy types to plant species
#' based on the morphophysiological CSR model by Novakovskiy et al. (2016), as well as performing
#' the required trait transformations for the model.
#'
#' @param data A data frame containing species names and trait data. Must include (LFW optional in place of LDMC):
#' \describe{
#'   \item{CH}{Canopy height (mm)}
#'   \item{LDMC}{Leaf dry matter content (\%)}
#'   \item{FP}{Flowering period (months)}
#'   \item{LS}{Lateral spread (categorical scale: 1–6, based on Hodgson et al. 1999)}
#'   \item{LDW}{Leaf dry weight (mg)}
#'   \item{LFW}{Leaf fresh weight(mg)}
#'   \item{PN}{Net photosynthesis (mg CO₂/g dry weight per hour)}
#'   \item{RD}{Dark respiration rate (same units as PN)}
#'   \item{LNC}{Leaf nitrogen content (mg/g)}
#'   \item{LCC}{Leaf carbon concentration (mg/g)}
#' }
#' @param calcLDMC Logical. Whether to calculate LDMC from LDW and LFW. Default is FALSE.
#'
#' @return A data frame with transformed traits, CSR percentages (C%, S%, R%), strategy classification and optional trait calculations.
#'
#' @import dplyr
#' @export
#'
#' @examples
#' # morphoPhys(exampleData)

morphoPhys <- function(data, calcLDMC = FALSE) {

  # column renaming helper function
  renameColumnsMorphophys <- function(data) {
    # making a list for names = required columns, values are regex patterns that might appear in a data set
    # list allows for multiple regexes
    renameDict <- list(
      # e.g. 'species' is the target name, telling the function that the values are equivalent
      # ^ and $ get exact matches so any word with sp in it will not match, '[._ ]?' matches users choice of spacing
      "species" = c("^species$", "^sp\\.$", "^sp$", "^taxon$", "^species[._ ]?name$", "^taxon[._ ]?name$"),
      "CH" = c("^canopy[._ ]?height$", "^CH$"),
      "LDMC" = c("^leaf[._ ]?dry[._ ]?matter[._ ]?content$", "^LDMC$"),
      "FP" = c("^flowering[._ ]?period$", "^FP$"),
      "LS" = c("^lateral[._ ]?spread$", "^LS$"),
      "LDW" = c("^leaf[._ ]?dry[._ ]?weight$", "^LDW$"),
      "LFW" = c("^leaf[._ ]?fresh[._ ]?weight$", "^LFW$"),
      "PN" = c("^net[._ ]?photosynthesis$", "^PN$"),
      "RD" = c("^dark[._ ]?respiration[._ ]?rate$", "^respiration[._ ]?rate$", "^RD$"),
      "LNC" = c("^leaf[._ ]?nitrogen[._ ]?content$", "^LNC$"),
      "LCC" = c("^leaf[._ ]?carbon[._ ]?concentration$", "^LCC$")
    )
    # for each user column loop through each name in dictionary (renameDict)
    for (colName in names(renameDict)) {
      # use grep() to find and return user columns matching a single regex patterns
      # patterns are taken from renameDict[[colName]] and collapsed into one regex string with paste(..., collapse = "|") to work with grep
      matchedCols <- grep(paste(renameDict[[colName]], collapse = "|"), names(data), ignore.case = TRUE, value = TRUE)
      # if at least one column matches...
      if (length(matchedCols) > 0) {
        # rename the matching column to the name in the dictionary list
        names(data)[names(data) %in% matchedCols] <- colName
      }
    }

    return(data)
  }

  # rename columns
  data <- renameColumnsMorphophys(data)

  # check for species column
  if (!"species" %in% names(data)) {
    stop("Data must contain a 'species' column.")
  }

  # check for required columns
  if (calcLDMC) {
    requiredCols <- c("species", "CH", "FP", "LS", "LDW", "PN", "RD", "LNC", "LCC", "LFW")
  } else {
    requiredCols <- c("species", "CH", "LDMC", "FP", "LS", "LDW", "PN", "RD", "LNC", "LCC")
  }

  # check column presence
  if (!all(requiredCols %in% names(data))) {
    stop(paste("Data must contain:", paste(requiredCols, collapse = ", ")))
  }

  # remove rows with missing values
  # pipeline to filter user data through - if any required columns contain na
  missingRows <- data %>% filter(if_any(all_of(requiredCols), ~ is.na(.)))
  if (nrow(missingRows) > 0) {
    # paste species names of rows with missing values into a vector then collapse into a string to be readable
    # e.g. (from c("species1",... to species1,...))
    removedSpecies <- paste(unique(missingRows$species), collapse = ", ")
    # display warning
    warning(sprintf(
      "%d row(s) removed due to missing trait values. affected species: %s",
      nrow(missingRows),
      removedSpecies
    ))
  }
  # anti_join to keep rows that do not match defined missingrows based on species column
  data <- anti_join(data, missingRows, by = "species")

  # ldmc calculation
  if (calcLDMC) {
    # pipeline to add new LDMC column via mutate
    data <- data %>% mutate(LDMC = (LDW * 100) / LFW)
  }

  # trait transformations
  calculations <- data %>%
    mutate(
      chTransformed   = ifelse(CH > 0, log(CH), NA),
      ldmcTransformed = ifelse(LDMC > 0, sqrt(LDMC), NA),
      fpTransformed   = FP,
      lsTransformed   = LS,
      ldwTransformed  = ifelse(LDW > 0, log(LDW) + 3, NA),
      pnTransformed   = ifelse(PN > 0, sqrt(PN), NA),
      rdTransformed   = ifelse(RD > 0, sqrt(RD), NA),
      lncTransformed  = ifelse(LNC > 0, sqrt(LNC), NA),
      lccTransformed  = ifelse(LCC > 0, sqrt(LCC), NA)
    )

  # pca score calculation
  calculations <- calculations %>%
    mutate(
      pca1 = 3.52173 + (0.21179 * chTransformed) - (0.26218 * ldmcTransformed) + (0.19528 * fpTransformed) -
        (0.04086 * lsTransformed) + (0.00836 * ldwTransformed) + (0.32013 * pnTransformed) +
        (1.00069 * rdTransformed) + (0.2593 * lncTransformed) - (0.38065 * lccTransformed),

      pca2 = -15.44964 + (0.58035 * chTransformed) + (0.42225 * ldmcTransformed) - (0.05119 * fpTransformed) +
        (0.21762 * lsTransformed) + (0.22002 * ldwTransformed) + (0.02368 * pnTransformed) +
        (0.35671 * rdTransformed) + (0.27781 * lncTransformed) + (0.28015 * lccTransformed)
    )

  # project onto csr axes
  calculations <- calculations %>%
    mutate(
      cScore = ((pca1 * 0.032) + (pca2 * 1.14)) / 1.14,
      sScore = ((pca1 * -0.842) + (pca2 * -0.678)) / 1.081,
      rScore = ((pca1 * 1.091) + (pca2 * -0.661)) / 1.276
    )

  # convert scores to percentages
  calculations <- calculations %>%
    mutate(
      # offset raw scores into positive range and convert to %
      denom = cScore + sScore + rScore + 6,
      cRaw = 100 * (cScore + 2) / denom,
      sRaw = 100 * (sScore + 2) / denom,
      rRaw = 100 * (rScore + 2) / denom,

      # clip any negative results to zero
      # pmax takes largest value, i.e. if cRaw = < 0, take 0
      cClipped = pmax(0, cRaw),
      sClipped = pmax(0, sRaw),
      rClipped = pmax(0, rRaw),

      # calculate how much total % was lost due to clipping
      deficit = 100 - (cClipped + sClipped + rClipped),

      # count how many axes had positive values (to share deficit across)
      positiveCount = (cClipped > 0) + (sClipped > 0) + (rClipped > 0),

      # redistribute lost percentage evenly to non-zero axes
      # e.g. if cClipped is bigger than 0, + the deficit divided by the amount of columns that aren't being clipped to be even.
      cPercent = cClipped + ifelse(cClipped > 0, deficit / positiveCount, 0),
      sPercent = sClipped + ifelse(sClipped > 0, deficit / positiveCount, 0),
      rPercent = rClipped + ifelse(rClipped > 0, deficit / positiveCount, 0)
    ) %>%
    # select, - means exclude that, so select the column that isn't raw, clipped etc. leaving cPercent, sPercent and rPercent, removes all other columns
    select(-denom, -ends_with("Raw"), -ends_with("Clipped"), -deficit, -positiveCount)

  # strategy classification
  # create a data frame to assign CSR value Strategy classes
  csrReference <- data.frame(
    strategy = c("C", "C/CR", "C/CS", "CR", "C/CSR", "CS", "CR/CSR", "CS/CSR", "R/CR",
                 "CSR", "S/CS", "R/CSR", "S/CSR", "R", "SR/CSR", "S", "R/SR", "S/SR", "SR"),
    cRef = c(2, 1, 1, 0, 1, 0, 0, 0, -1, 0, -1, -1, -1, -2, -1, -2, -2, -2, -2),
    sRef = c(-2, -2, -1, -2, -1, 0, -1, 0, -2, 0, 1, -1, 1, -2, 0, 2, -1, 1, 0),
    rRef = c(-2, -1, -2, 0, -1, -2, 0, -1, 1, 0, -2, 1, -1, 2, 0, -2, 1, -1, 0)
  )
  # classify function for c s and r scores (in use c = cScore etc.)
  classifySpecies <- function(c, s, r) {
    # distance between c s and r scores (coordinates) and reference c s and r from referencde data frame
    distances <- sqrt((csrReference$cRef - c)^2 + (csrReference$sRef - s)^2 + (csrReference$rRef - r)^2)
    # which.min finds row number with smallest variance, variance$strategy[] - square brackets get the value from that row
    return(csrReference$strategy[which.min(distances)])
  }

  calculations <- calculations %>%
    # apply by row instead of whole column
    rowwise() %>%
    mutate(strategyClass = classifySpecies(cScore, sScore, rScore)) %>%
    # ungroup from rowwise back to column based
    ungroup()

  # define the output columns
  outputCols <- c("cScore", "sScore", "rScore", "cPercent", "sPercent", "rPercent", "strategyClass")
  if (calcLDMC) outputCols <- c("LDMC", outputCols)

  # combine calculations columns with user data via cbind
  finalResult <- cbind(
    data,
    calculations %>% dplyr::select(dplyr::any_of(outputCols))
  )

  return(finalResult)
}
