## Supplementary material for "CSR Calculator: An R package and Shiny application for assigning plant ecological strategies using trait data": R code text files used in the CSRcalculator R package and shiny application.: strateFy.txt

#' Calculate CSR Scores and Assign a CSR Strategy Using the StrateFy Model (Pierce et al. 2017)
#'
#' This function calculates and assigns CSR strategy types to plant species
#' based on the StrateFy model, as well as calculating required leaf traits for the model.
#'
#' @param data A data frame containing species names and trait data. Must include either:
#' \itemize{
#'   \item \strong{LA}, \strong{LFW}, and \strong{LDW}: Leaf area (mm²), leaf fresh weight (mg), and leaf dry weight (mg)
#'   \item \strong{LA}, \strong{SLA}, and \strong{LDMC}: Leaf area (mm²), specific leaf area (mm²/mg), and leaf dry matter content (%)
#' }
#' @param calcSLA Logical. Whether to calculate SLA if not provided. Default is TRUE.
#' @param calcLDMC Logical. Whether to calculate LDMC if not provided. Default is TRUE.
#'
#' @return A data frame with calculated CSR percentages (C%, S%, R%), strategy classification, and optionally SLA, LDMC, and succulence index.
#'
#' @import dplyr
#' @export
#'
#' @examples
#' # strateFy(exampleData)

strateFy <- function(data, calcSLA = TRUE, calcLDMC = TRUE) {

  # column renaming helper function
  renameColumnsStratefy <- function(data) {
    # making a list for names = required columns, values are regex patterns that might appear in a data set
    # list allows for multiple regexes
    renameDict <- list(
      # e.g. 'species' is the target name, telling the function that the values are equivalent
      # ^ and $ get exact matches so any word with sp in it will not match, '[._ ]?' matches users choice of spacing
      "species" = c("^species$", "^sp\\.$", "^sp$", "^taxon$", "^species[._ ]?name$", "^taxon[._ ]?name$"),
      "LA" = c("^leaf[._ ]?area$", "^LA$"),
      "LFW" = c("^leaf[._ ]?fresh[._ ]?weight$", "^LFW$"),
      "LDW" = c("^leaf[._ ]?dry[._ ]?weight$", "^LDW$"),
      "SLA" = c("^specific[._ ]?leaf[._ ]?area$", "^SLA$"),
      "LDMC" = c("^leaf[._ ]?dry[._ ]?matter[._ ]?content$", "^LDMC$")
    )
# for each user column loop through each name in dictionary (renameDict)
    for (colName in names(renameDict)) {
      # use grep() to find and return user columns matching a single regex patterns
      # patterns are taken from renameDict[[colName]] and collapsed into one regex string with paste(..., collapse = "|") to work with grep
      matchedCols <- grep(paste(renameDict[[colName]], collapse = "|"), names(data), ignore.case = TRUE, value = TRUE)
      # if at least one column matches...
      if (length(matchedCols) > 0) {
        # rename the matching column to the name in the dictionary list
        names(data)[names(data) %in% matchedCols] <- colName
      }
    }

    return(data)
  }

  # rename columns
  data <- renameColumnsStratefy(data)

  # check for species column
  if (!"species" %in% names(data)) {
    stop("Data must contain a 'species' column")
  }

  # defining the required columns
  if (calcSLA && calcLDMC) {
    requiredCols <- c("species", "LA", "LFW", "LDW")
  } else if (calcSLA) {
    requiredCols <- c("species", "LA", "LDW", "LDMC")
  } else if (calcLDMC) {
    requiredCols <- c("species", "LA", "LFW", "SLA")
  } else {
    requiredCols <- c("species", "LA", "SLA", "LDMC")
  }

  # check for the required columns presence
  if (!all(requiredCols %in% names(data))) {
    stop(paste("Data must contain:", paste(requiredCols, collapse = ", ")))
  }

  # remove rows with missing values
  # pipeline to filter user data through - if any required columns contain na
  missingRows <- data %>% filter(if_any(all_of(requiredCols), ~ is.na(.)))
  if (nrow(missingRows) > 0) {
    # paste species names of rows with missing values into a vector then collapse into a string to be readable
    # e.g. (from c("species1",... to species1,...))
    removedSpecies <- paste(unique(missingRows$species), collapse = ", ")
    # display warning
    warning(sprintf(
      "%d row(s) removed due to missing trait values. affected species: %s",
      nrow(missingRows),
      removedSpecies
    ))
  }
  # anti_join to keep rows that do not match defined missingrows based on species column
  data <- anti_join(data, missingRows, by = "species")

  # sla calculation
  if (calcSLA) {
    # pipeline to add new SLA column via mutate
    data <- data %>% mutate(SLA = LA / LDW)
  }

  # ldmc calculation
  if (calcLDMC) {
    data <- data %>%
      mutate(
        succulenceIndex = (LFW - LDW) / (LA / 10),
        LDMC = ifelse(
          succulenceIndex > 5,
          (100 - (LDW * 100) / LFW),
          (LDW * 100) / LFW
        )
      )
  }

  # define bounds and translation constants
  maxCDim <- 57.376; minCDim <- 0
  maxSDim <- 5.792; minSDim <- -0.756
  maxRDim <- 1.10795515716546; minRDim <- -11.3467682227961
  posTranslationCoefC <- abs(minCDim)
  posTranslationCoefS <- abs(minSDim)
  posTranslationCoefR <- abs(minRDim)

  # trait transformations and csr score calculations
  calculations <- data %>%
    mutate(
      # c axis calculation
      sqrtMaxLA = sqrt(LA / 894205) * 100,
      pca2C = -0.8678 + 1.6464 * sqrtMaxLA,
      # pmin and pmax return the smaller or larger of the two values (e.g. for pmax if pca2c is smaller that mincdim return the mincdim)
      pca2C = pmin(pmax(pca2C, minCDim), maxCDim),
      cTrans = pca2C + posTranslationCoefC,
      cRange = maxCDim + posTranslationCoefC,
      propTotalVariabilityC = (cTrans / cRange) * 100,

      # s axis calculation
      logitLdmc = log((LDMC / 100) / (1 - (LDMC / 100))),
      # exp = e raised to the power of (x), e.g. e^(-0.2328 * logitLdmc)
      pca1S = 1.3369 + 0.000010019 * (1 - exp(-2.2303e-12 * logitLdmc)) + 4.5835 * (1 - exp(-0.2328 * logitLdmc)),
      pca1S = pmin(pmax(pca1S, minSDim), maxSDim),
      sTrans = pca1S + posTranslationCoefS,
      sRange = maxSDim + posTranslationCoefS,
      propTotalVariabilityS = (sTrans / sRange) * 100,

      # r axis calculation
      logSla = log(SLA),
      pca1R = -57.5924 + 62.6802 * exp(-0.0288 * logSla),
      pca1R = pmin(pmax(pca1R, minRDim), maxRDim),
      rTrans = pca1R + posTranslationCoefR,
      rRange = maxRDim + posTranslationCoefR,
      propTotalVariabilityR = 100 - (rTrans / rRange * 100),

      # convert to percentages
      percentageConversionCoeff = 100 / (propTotalVariabilityC + propTotalVariabilityS + propTotalVariabilityR),
      cPercent = propTotalVariabilityC * percentageConversionCoeff,
      sPercent = propTotalVariabilityS * percentageConversionCoeff,
      rPercent = propTotalVariabilityR * percentageConversionCoeff
    )

  # strategy classification
  # create a data frame to assign CSR value Strategy classes
  zoneValues <- data.frame(
    strategy = c("C", "C/CR", "C/CS", "CR", "C/CSR", "CS", "CR/CSR", "CS/CSR", "R/CR",
                 "CSR", "S/CS", "R/CSR", "S/CSR", "R", "SR/CSR", "S", "R/SR", "S/SR", "SR"),
    C = c(90, 73, 73, 48, 54, 48, 42, 42, 23, 33, 23, 23, 23, 5, 17, 5, 5, 5, 5),
    S = c(5, 5, 23, 5, 23, 48, 17, 42, 5, 33, 73, 23, 54, 5, 42, 90, 23, 73, 48),
    R = c(5, 23, 5, 48, 23, 5, 42, 17, 73, 33, 5, 54, 23, 90, 42, 5, 73, 23, 48)
  )

  # pipeline to calculate closest strategy for each row of the users data set based on C S and R %s
  calculations <- calculations %>%
    # apply by row instead of whole column
    rowwise() %>%
    # create class column
    mutate(
      strategyClass = {
        # temp data frame called variances based on earlier zonevalues data frame with predefined strategies
        variances <- zoneValues %>%
          # add column to temp dataframe calculating variance between CSR %s and predefined values
          mutate(
            variance = (cPercent - C)^2 +
              (sPercent - S)^2 +
              (rPercent - R)^2
          )
        # which.min finds row number with smallest variance, variance$strategy[] - square brackets get the value from that row
        variances$strategy[which.min(variances$variance)]
      }
    ) %>%
    # ungroup from rowwise back to column based
    ungroup()

  # define the output columns
  outputCols <- c("cPercent", "sPercent", "rPercent", "strategyClass")
  if (calcSLA) outputCols <- c("SLA", outputCols)
  if (calcLDMC) outputCols <- c("LDMC", "succulenceIndex", outputCols)

  # combine calculations columns with user data frame via cbind
  finalResult <- cbind(
    data,
    # select any of the outputcols, as they will only exist if conditionals are met
    calculations %>% dplyr::select(dplyr::any_of(outputCols))
  )

  return(finalResult)
}
