## Supplementary material for "CSR Calculator: An R package and Shiny application for assigning plant ecological strategies using trait data": Equations and methods used for each model function in the CSRcalculator R package.

StrateFy (Pierce et al. 2017)

Traits:

- LA: Leaf area (mm²)
- LFW: Leaf fresh weight (mg)
- LDW: Leaf dry weight (mg)

Trait transformations:

- SLA = LA / LDW
- succulenceIndex = (LFW − LDW) / (LA / 10)
- LDMC =
   If succulenceIndex > 5: 100 − (LDW × 100 / LFW)
   Else: (LDW × 100) / LFW

CSR axis scores:

C-axis:

- sqrtMaxLA = sqrt(LA / 894205) × 100
- pca2C = −0.8678 + 1.6464 × sqrtMaxLA
- Clamp pca2C between model limits: minCdimension = 0, maxCdimension = 57.376
- Translated C-axis value = pca2C + abs(minCdimension)
- Total C-axis range after translation = maxCdimension + abs(minCdimension)
- Proportional C-axis score = (Translated C-axis value / Total C-axis range) × 100

S-axis:

- logitLDMC = log((LDMC / 100) / (1 − LDMC / 100))
- pca1S = 1.3369 + 0.000010019 × (1 − exp(−2.2303 × 10⁻¹² × logitLDMC)) + 4.5835 × (1 − exp(−0.2328 × logitLDMC))
- Clamp pca1S between model limits: minSdimension = −0.756, maxSdimension = 5.792
- Translated S-axis value = pca1S + abs(minSdimension)
- Total S-axis range after translation = maxSdimension + abs(minSdimension)
- Proportional S-axis score = (Translated S-axis value / Total S-axis range) × 100

R-axis:

- logSLA = log(SLA)
- pca1R = −57.5924 + 62.6802 × exp(−0.0288 × logSLA)
- Clamp pca1R between model limits: minRdimension = −11.347, maxRdimension = 1.108
- Translated R-axis value = pca1R + abs(minRdimension)
- Total R-axis range after translation = maxRdimension + abs(minRdimension)
- Proportional R-axis score = 100 − ((Translated R-axis value / Total R-axis range) × 100)

Percentage conversion:

- conversionCoeff = 100 / (C + S + R)
- C% = C × conversionCoeff
- S% = S × conversionCoeff
- R% = R × conversionCoeff

Strategy assignment:

Each species is assigned to the nearest predefined strategy:

$$\text{strategy}=\min\left( \left( C_{\text{ref}}-C\% \right)^{2}+\left( S_{\text{ref}}-S\% \right)^{2}+\left( R_{\text{ref}}-R\% \right)^{2} \right)$$

Predefined strategies:

| Strategy | Cref | Sref | Rref |
| --- | --- | --- | --- |
| C | 90 | 5 | 5 |
| C/CR | 73 | 5 | 23 |
| C/CS | 73 | 23 | 5 |
| CR | 48 | 5 | 48 |
| C/CSR | 54 | 23 | 23 |
| CS | 48 | 48 | 5 |
| CR/CSR | 42 | 17 | 42 |
| CS/CSR | 42 | 42 | 17 |
| R/CR | 23 | 5 | 73 |
| CSR | 33 | 33 | 33 |
| S/CS | 23 | 73 | 5 |
| R/CSR | 23 | 23 | 54 |
| S/CSR | 23 | 54 | 23 |
| R | 5 | 5 | 90 |
| SR/CSR | 17 | 42 | 42 |
| S | 5 | 90 | 5 |
| R/SR | 5 | 5 | 90 |
| S/SR | 5 | 73 | 23 |
| SR | 5 | 48 | 48 |

Morpho-physiological (Novakovskiy et al. 2021)

Traits:

- CH: Canopy height (mm)
- LDMC: Leaf dry matter content (%)
- FP: Flowering period (months)
- LS: Lateral spread (categorical scale: 1–6, see Hodgson et al. (1999))
- LDW: Leaf dry weight (mg)
- LFW: Leaf fresh weight (mg)
- PN: Net photosynthesis (mg CO₂/g dry weight per hour)
- RD: Dark respiration rate (mg CO₂/g dry weight per hour)
- LNC: Leaf nitrogen content (mg/g)
- LCC: Leaf carbon concentration (mg/g)

Trait transformations:

- CH: log(CH)
- LDW: log(LDW) + 3
- LDMC, PN, RD, LNC, LCC: sqrt(value)

CSR axis scores:

PCA:

- pca1 = 3.52173 + 0.21179 × log(CH) − 0.26218 × sqrt(LDMC) + 0.19528 × FP − 0.04086 × LS + 0.00836 × log(LDW) + 0.32013 × sqrt(PN) + 1.00069 × sqrt(RD) + 0.2593 × sqrt(LNC) − 0.38065 × sqrt(LCC)
- pca2 = −15.44964 + 0.58035 × log(CH) + 0.42225 × sqrt(LDMC) − 0.05119 × FP + 0.21762 × LS + 0.22002 × log(LDW) + 0.02368 × sqrt(PN) + 0.35671 × sqrt(RD) + 0.27781 × sqrt(LNC) + 0.28015 × sqrt(LCC)

CSR projection:

- C = ((pca1 × 0.032) + (pca2 × 1.14)) / 1.14
- S = ((pca1 × −0.842) + (pca2 × −0.678)) / 1.081
- R = ((pca1 × 1.091) + (pca2 × −0.661)) / 1.276

Percentage conversion:

- C% = 100 × (C + 2) / (C + S + R + 6)
- S% = 100 × (S + 2) / (C + S + R + 6)
- R% = 100 × (R + 2) / (C + S + R + 6)

If any of these are negative, they are set to 0. The difference is redistributed evenly among the non-zero axes.

Strategy assignment:

Each species is assigned to the closest predefined strategy:

$$\text{strategy}=\min\left( \sqrt{\left( C_{\text{ref}}-C \right)^{2}+\left( S_{\text{ref}}-S \right)^{2}+\left( R_{\text{ref}}-R \right)^{2}} \right)$$

Predefined strategies:

| Strategy | Cref | Sref | Rref |
| --- | --- | --- | --- |
| C | 2 | -2 | -2 |
| C/CR | 1 | -2 | -1 |
| C/CS | 1 | -1 | -2 |
| CR | 0 | -2 | 0 |
| C/CSR | 1 | -1 | -1 |
| CS | 0 | 0 | -2 |
| CR/CSR | 0 | -1 | 0 |
| CS/CSR | 0 | 0 | -1 |
| R/CR | -1 | -2 | 1 |
| CSR | 0 | 0 | 0 |
| S/CS | -1 | 1 | -2 |
| R/CSR | -1 | -1 | 1 |
| S/CSR | -1 | 1 | -1 |
| R | -2 | -2 | 2 |
| SR/CSR | -1 | 0 | 0 |
| S | -2 | 2 | -2 |
| R/SR | -2 | -2 | 0 |
| S/SR | -2 | 1 | -1 |
| SR | -2 | 0 | 0 |

Hodgson et al. (1999)

Traits:

- CH: Canopy height (mm)
- LDMC: Leaf dry matter content (%)
- FP: Flowering period (months)
- LS: Lateral spread (categorical scale: 1–6, from Hodgson et al. 1999)
- LDW: Leaf dry weight (mg)
- SLA: Specific leaf area (mm²/mg)
- LFW: Leaf fresh weight (mg)
- LA: Leaf area (mm²)
- FS: Flowering start (1–6, where 1 = March or earlier, 6 = August or later)

Trait transformations:

- CH is converted into a six-point ordinal scale: CH > 999 = 6, CH > 599 = 5, CH > 299 = 4, CH > 99 = 3, CH > 49 = 2, Else = 1
- LDW = log(LDW) + 3
- LDMC and SLA = sqrt(value)

CSR axis scores:

Two separate sets of equations are used depending on whether the species is considered a grass.

- Grass version:
  - rawC = (0.141 × CH²) + (0.09061 × LS²)
  - rawS = 54.6 – (1.666 × chPr²) + (1.069 × LDMC²) – (2.732 × SLA²) + (1.722 × LS²)
  - rawR = (2.518 × FP) – (2.748 × LDW) + (5.37 × SLA)
- Non-grass version:
  - rawC = (0.09245 × CH²) + (0.05631 × LS²) + (0.01595 × LDW²)
  - rawS = −39.52 – (7.581 × CH) + (2.633 × LDMC²) – (0.351 × LDW²)
  - rawR = −(1.158 × LDMC²) + (3.137 × FP) + (3.145 × FS) – (0.0849 × LDW²) – (1.193 × SLA²) + (11.4 × SLA)

Score transformation:

- C = −2.5 + 0.839 × rawC
- S = −1.103 + 0.0474 × rawS (grass) or S = −1.249 + 0.0531 × rawS (non-grass)
- R = −2.5 + 0.119 × rawR

Scores are then rounded down to the nearest tenth after being clipped to the range –2.5 to 2.5.

Percentage conversion:

- C% = 100 × (C + 2.5) / (C + S + R + 7.5)
- S% = 100 × (S + 2.5) / (C + S + R + 7.5)
- R% = 100 × (R + 2.5) / (C + S + R + 7.5)

Strategy assignment:

Same as morpho-physiological.

Predefined strategies:

| Strategy | Cref | Sref | Rref |
| --- | --- | --- | --- |
| C | 2 | -2 | -2 |
| C/CR | 1 | -2 | -1 |
| C/SC | 1 | -1 | -2 |
| CR | 0 | -2 | 0 |
| C/CSR | 1 | -1 | -1 |
| SC | 0 | 0 | -2 |
| CR/CSR | 0 | -1 | 0 |
| SC/CSR | 0 | 0 | -1 |
| R/CR | -1 | -2 | 1 |
| CSR | 0 | 0 | 0 |
| S/SC | -1 | 1 | -2 |
| R/CSR | -1 | -1 | 1 |
| S/CSR | -1 | 1 | -1 |
| R | -2 | -2 | 2 |
| SR/CSR | -1 | 0 | 0 |
| S | -2 | 2 | -2 |
| R/SR | -2 | -2 | 0 |
| S/SR | -2 | 1 | -1 |
| SR | -2 | 0 | 0 |
