## Supplementary material for "CSR Calculator: An R package and Shiny application for assigning plant ecological strategies using trait data": Comparison of CSR scores produced by model sources and the CSRcalculator application.

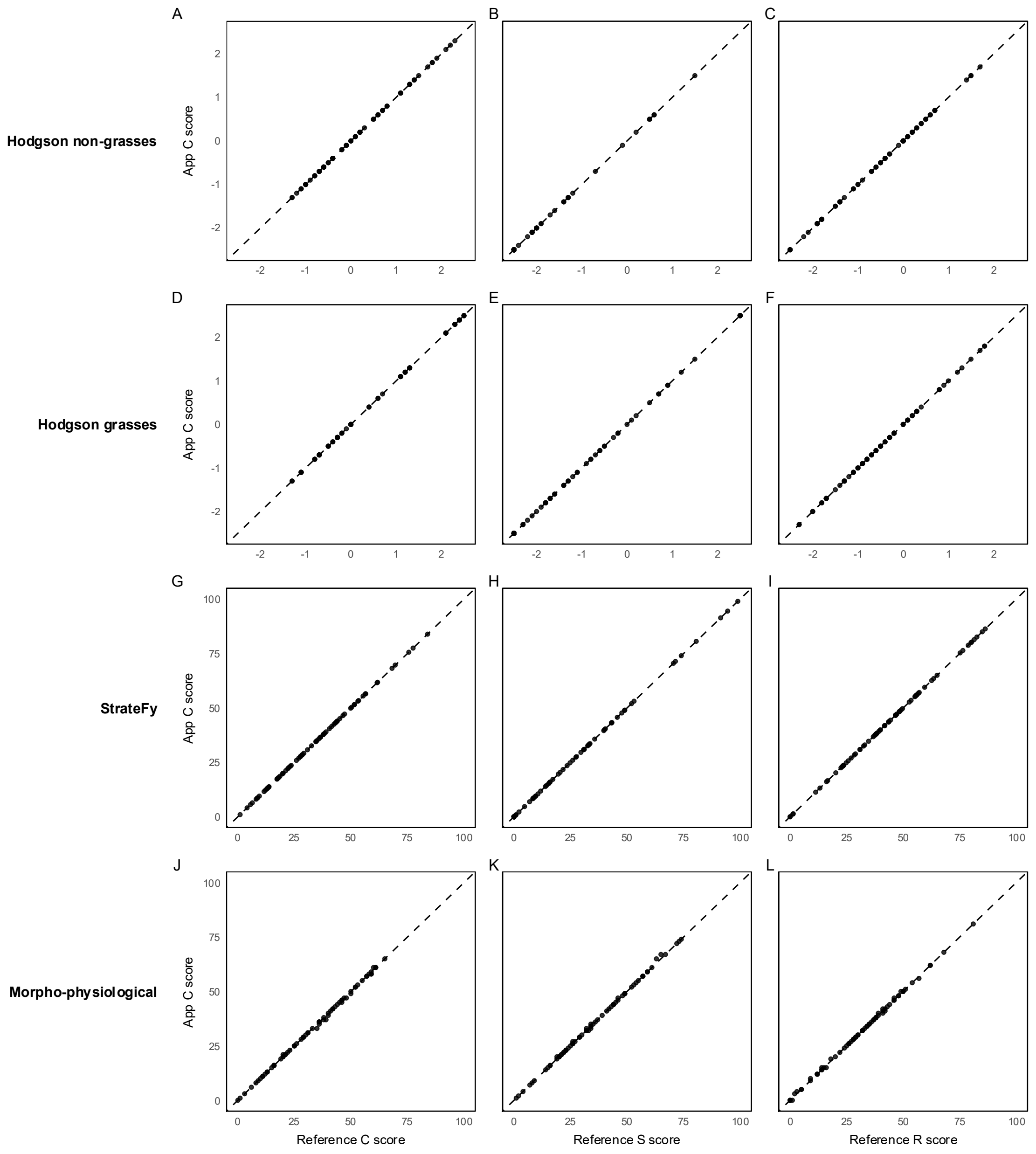


Figure S1. Comparison between C (A, D, G, J), S (B, E, H, K), R (C, F, I, L) score outputs from the *CSRcalculator* tools (Application) and the respective model reference: StrateFy (Pierce et al. 2017) and Hodgson et al. (1999) excel tools, morphophysiological model output from Novakovskiy et al. (2021) supplementary data. Trait data used is from Novakovskiy et al. (2021) supplementary data.
